# Real-Time Closed-Loop Feedback System For Mouse Mesoscale Cortical Signal And Movement Control: CLoPy

**DOI:** 10.1101/2024.11.02.619716

**Authors:** Pankaj K. Gupta, Timothy H. Murphy

**Author notes:** Correspondence to: Timothy H Murphy, Address: Djavad Mowafaghian Centre for Brain Health (DMCBH), 2215 Wesbrook Mall, Vancouver, BC V6T 1Z3, Canada.

## Abstract

Increasingly, experiments designed to provide practical perturbations to circuits or behavior are required for hypothesis testing in various disciplines ranging from motor learning to recovery after injury. We present the implementation and efficacy of an open-source closed-loop neurofeedback (CLNF) and closed-loop movement feedback (CLMF) system. In CLNF, we measure mm-scale cortical mesoscale activity with GCaMP6s and provide graded auditory feedback (within ∼63 ms) based on changes in dorsal-cortical activation within regions of interest (ROI) and with a specified rule. Single or dual ROIs (ROI1, ROI2) on the dorsal cortical map were selected as targets. Both motor and sensory regions supported closed-loop training in male and female mice. Mice modulated activity in rule-specific target cortical ROIs to get increasing rewards over days (RM ANOVA p=2.83e-5) and adapted to changes in ROI rules (RM ANOVA p=8.3e-10, Table 4 for different rule changes). In CLMF, feedback (within ∼67 ms) was based on tracking a specified body movement, and rewards were generated when the behavior reached a threshold. For movement training, the group that received graded auditory feedback performed significantly better (RM-ANOVA p=9.6e-7) than a control group (RM-ANOVA p=0.49) within four training days. Additionally, mice can learn a change in task rule from left forelimb to right forelimb within a day, after a brief performance drop on day 5. Offline analysis of neural data and behavioral tracking revealed changes in the overall distribution of Ca2+ fluorescence values in CLNF and body-part speed values in CLMF experiments. Increased CLMF performance was accompanied by a decrease in task latency and cortical ΔF/F_0_ amplitude during the task, indicating lower cortical activation as the task gets more familiar.

## Introduction

In neuroscience, brain activity and behavior are often studied as separate entities that do not interact in real time. Assessments are typically performed post hoc, and experimental designs rarely incorporate dynamic feedback based on ongoing neural activity. In contrast, closed-loop systems provide a framework where real-time interaction occurs between the brain, subject, and goal-directed outcomes, enabling precise modulation of neural and behavioral dynamics (Clancy et al. 2014). First introduced in the pioneering work of (Fetz 1969), closed-loop feedback has since demonstrated significant potential in understanding and modulating brain-behavior relationships. There has been recent interest in its application to rodent models (Clancy and Mrsic-Flogel 2021; Clancy et al. 2014; Prsa, Galiñanes, and Huber 2017; Srinivasan et al. 2018; Knudsen and Wallis 2020; Paz et al. 2013; Ching et al. 2013; Rosin et al. 2011; Widge and Moritz 2014; Sun et al. 2022). Some related studies have demonstrated the feasibility of closed-loop feedback in rodents, including hippocampal electrical feedback to disrupt memory consolidation (Girardeau et al. 2009), optogenetic perturbations of somatosensory circuits during behavior (O’Connor et al. 2013), and more recent advances employing targeted optogenetic interventions to guide behavior (Abbasi et al. 2023). Opportunities also exist in extending real time pose classification (Forys et al. 2020; Kane et al. 2020) and movement perturbation (M. W. Mathis, Mathis, and Uchida 2017) to shape aspects of an animal’s motor repertoire. This gap is due, in part, to technical challenges such as the need for low-latency data acquisition and processing, compact and portable hardware, and seamless integration with animal behavior. Moreover, the inherent complexity of rodent neurophysiology and behavior (e.g., undesired movements, variability in neural signals, and behavioral metrics) further complicates the implementation of closed-loop paradigms.

Closed-loop systems are particularly promising for addressing critical questions in systems neuroscience, such as how brain activity dynamically adapts to feedback or how neural circuits reorganize during motor learning and recovery. Recent advancements, such as genetically encoded calcium indicators and mesoscopic imaging, have opened new avenues for studying these phenomena (Clancy and Mrsic-Flogel 2021; Clancy et al. 2014; Prsa, Galiñanes, and Huber 2017). However, many of the available closed-loop systems remain technically complex, expensive, or inaccessible to labs with limited resources. Additionally, most existing platforms focus on narrowly defined use cases, limiting their adaptability to diverse experimental paradigms. Nevertheless, recent work has demonstrated behavioral control through feedback, such as in (M. W. Mathis, Mathis, and Uchida 2017), where mice learned to pull a joystick to a target and adapt this motion under force perturbations.

To address these gaps, we present CLoPy - a flexible, cost-effective, and open-source Python-based platform for closed-loop feedback experiments in neuroscience. CLoPy integrates real-time data acquisition, processing, and feedback delivery, providing researchers with a robust tool to study dynamic brain-behavior interactions in head-fixed mice. The platform supports two modes of graded feedback: (1) Closed-Loop Neurofeedback (CLNF), which derives feedback from neuronal activity, and (2) Closed-Loop Movement Feedback (CLMF), which bases feedback on observed body movements. By combining genetically encoded calcium imaging with behavioral tracking, CLoPy enables researchers to explore questions such as how cortical dynamics contribute to motor learning or how feedback modulates neural plasticity in neurological models.

Crucially, CLoPy is designed with accessibility and versatility in mind. The platform utilizes widely available hardware, including Raspberry Pi and Nvidia Jetson, and leverages Python-based open-source software libraries. These features make CLoPy an ideal resource for the broader open-source neuroscience community, particularly for labs seeking to implement closed-loop systems without substantial financial or technical barriers. Furthermore, CLoPy’s modular design allows researchers to adapt it for a wide range of experimental paradigms, from studying cortical reorganization during motor learning to investigating sensory feedback mechanisms. By framing CLoPy within the context of accessible, open-source neuroscience tools, we aim to broaden its impact and utility across the systems neuroscience community.

In this study, we demonstrate the implementation and efficacy of CLoPy in head-fixed mice (Figure 1), showcasing its ability to deliver precise, low-latency feedback. We provide all software, hardware schematics, and protocols necessary for its adaptation to various experimental scenarios, positioning CLoPy as a valuable tool for advancing closed-loop neuroscience research.

**Figure 1.**
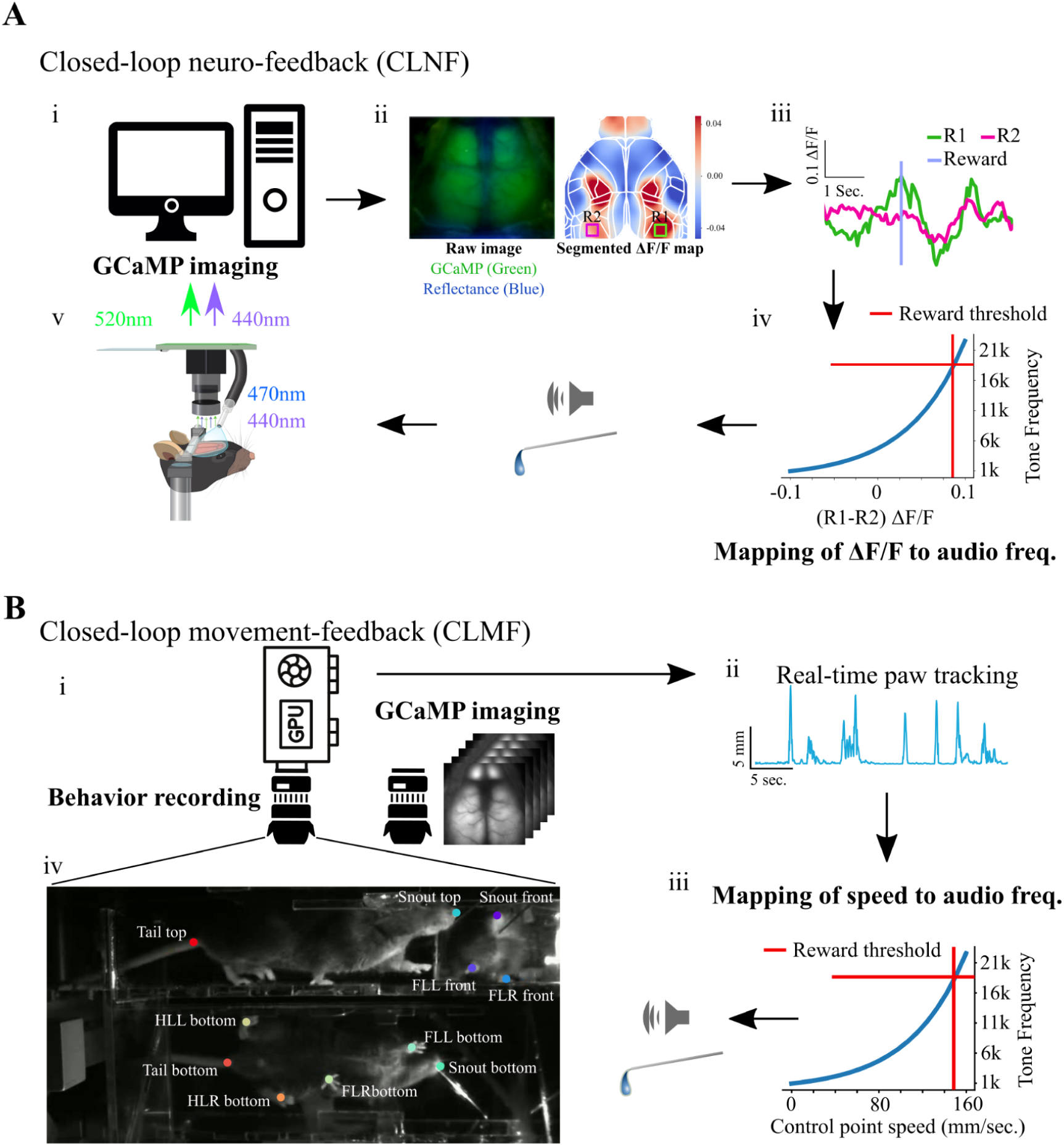
Setup of real-time feedback for GCaMP6 cortical activity and movements. A) GCaMP-based Closed-loop Feedback and Reward System: Mice expressing GCaMP6, with surgically implanted transcranial windows and a head-bar, were positioned beneath the imaging camera, with GCaMP6 excitation light at 470 nm. A secondary wavelength of 440 nm was used for continuous reflectance signals to measure hemodynamic changes, which were then applied to correct the fluorescence signal. The RGB camera was equipped with bandpass filters that allowed only 520 nm epifluorescence and 440 nm reflectance to be simultaneously collected. i) Mice with transcranial windows were head-fixed beneath an imaging camera, with the cortical window illuminated using 440 nm (for reflectance signal used for hemodynamic correction) and 470 nm (for GCaMP excitation) light. Epifluorescence at 520 nm and reflectance at 440 nm were captured at 15 fps using a bandpass filter, integrated within the cortical imaging system. ii) The captured images were simultaneously saved and processed to compute ΔF/F_0_ in real-time using a Raspberry Pi 4B model. Pre-selected regions of interest (ROIs) were continuously monitored, and rule-specific activation was calculated based on the ΔF/F_0_ signal. The left panel displayed wide-field cortical GCaMP6 fluorescence (green) and reflectance (blue), while the right panel showed the real-time calculated and corrected ΔF/F_0_ map, generated using a moving average of the captured images. The target ROIs were marked as green (R1) and red (R2), although a single ROI could also be selected for monitoring. iii) For example, as shown on the ΔF/F_0_ map, ROIs R1 and R2 were continuously monitored, and the average activity across these ROIs was calculated. iv) When the task rule was defined as “R1 - R2 > threshold”, the difference between R1 and R2 activities was mapped to a non-linear function that generated graded audio tone frequencies (ranging from 1 kHz to 22 kHz), as illustrated in the figure. Task rules could be modified within the setup on any given day, and the corresponding activation levels were automatically mapped to the audio frequencies. The “threshold”, expressed in ΔF/F_0_ units as a percent change, was adjustable based on the experimental design. v) Upon reaching the rule-specific threshold for activity, in addition to the increase in audio tone frequency, a water reward was delivered to the head-fixed mouse. B) Closed-loop Behavior Feedback and Reward Setup: A specialized transparent head-fixation chamber was custom-designed using 3mm-thick plexiglass material (3D model available on GitHub link https://github.com/pankajkgupta/clopy) to enable multi-view behavioral imaging and real-time tracking of body parts. The rectangular chamber was equipped with two strategically positioned mirrors—one at the bottom, angled at 45 degrees, and one at the front, angled at 30 degrees—facilitating multi-view imaging of the head-fixed mouse with a single camera. i) A Dalsa CCD camera was connected to a PC for widefield cortical imaging during the session. An additional camera connected to a PC with a GPU was used to capture the behavior video stream of the headfixed mouse. ii) Video frames of the behavior were processed in real-time on a GPU (Nvidia Jetson Orin), which tracked the body parts using a custom trained DeepLabCut-Live(Kane et al. 2020) model. iii) Auditory feedback was provided using a non-linear function mapping paw speeds to corresponding audio tone frequencies. iv) The head-fixed mouse, positioned in the transparent chamber, was able to freely move its body parts, while its behavior was continuously recorded. This setup allowed for the capture of three distinct views of the mouse—side, front, and bottom profiles—and enabled the real-time tracking of multiple body parts, including the snout, left and right forelimbs, left and right hindlimbs, and the base of the tail.

**Figure 2:**
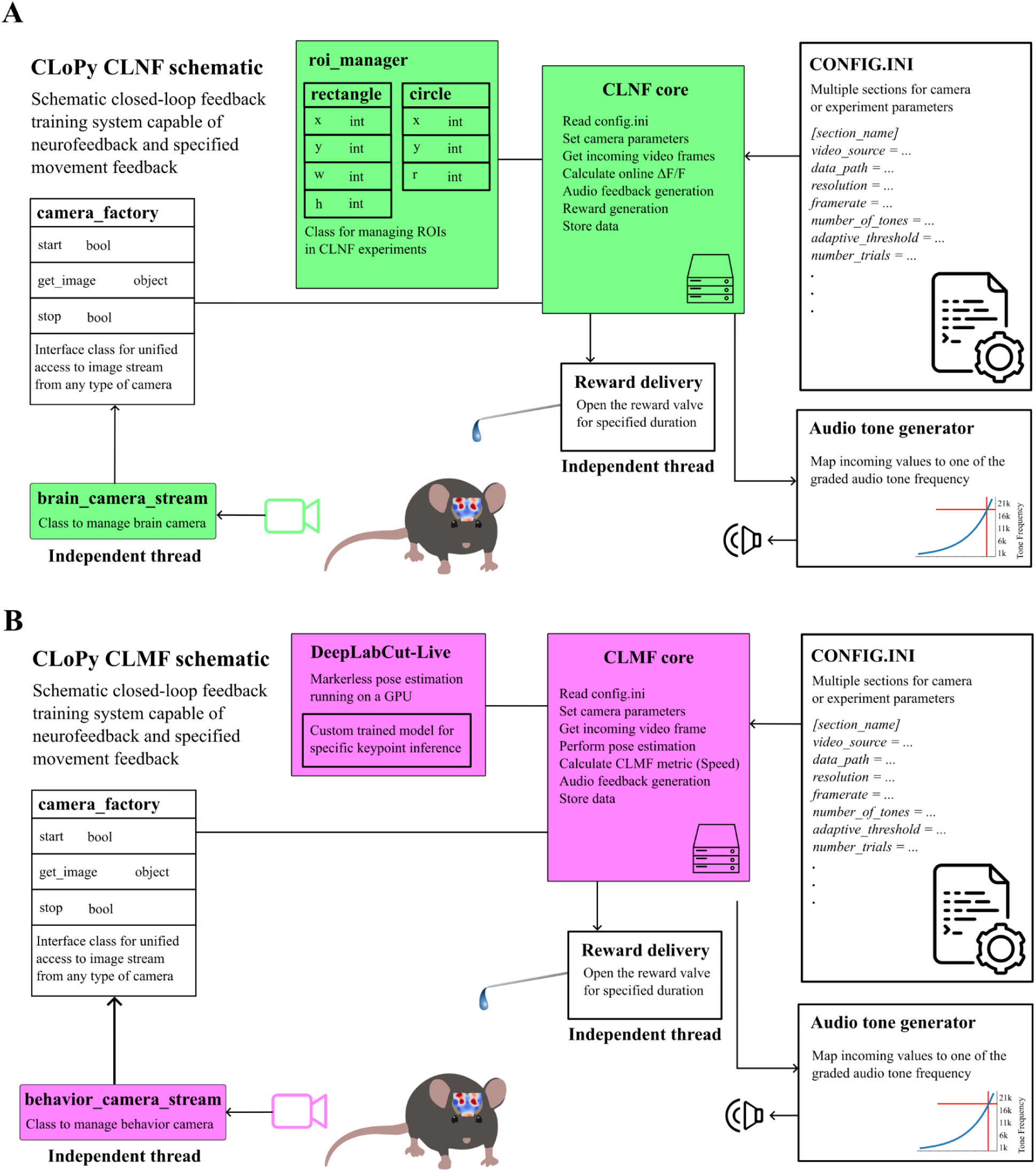
Schematic of the closed-loop feedback training system (CLoPy) for neurofeedback and specified movement feedback. A) The components of CLoPy CLNF system are presented in a block diagram. Modular components such as the configuration file, camera factory, audio tone generator, and reward delivery system are displayed and are utilized by both the Closed-Loop Neurofeedback (CLNF) and Closed-Loop Movement Feedback (CLMF, in B) systems. The configuration file (config.ini) stored all configurable parameters of the system, including camera settings, feedback parameters, reward thresholds, number of trials, and the duration of trial and rest periods, under an experiment-specific section. The camera factory was an abstract class that provided a unified interface for a programmable camera to the CLNF and CLMF(in B) systems. This abstraction allowed the core system to remain independent of the specific camera libraries required for image streaming. Camera-specific routines were implemented in separate “brain_camera_stream” and “behavior_camera_stream” classes, which inherited functions from the “camera_factory” superclass and ran in independent thread processes. The Region of Interest (ROI) manager was used by the CLNF core to maintain a list of ROIs, as well as routines to perform rule-specific operations on them, as specified in config.ini. An ROI could be defined as a rectangle (with the upper-left corner coordinates, height, and width) or as a circle (with center coordinates and radius). The audio tone generator mapped the target activity (fluorescence signal in CLNF) to graded audio tone frequencies. It generated audio signals at 44.1 kHz sampling based on the specified frequency and sent the signal to the audio output. Reward delivery was controlled by opening a solenoid valve for a specified duration, which was managed in a separate process thread. The CLNF core was the main program responsible for running the CLNF system. It utilized config.ini, the camera factory, the ROI manager, and integrated the audio tone generator and reward delivery functions. The system also saved the recorded data and configuration parameters with unique identifiers. B) The components of CLoPy CLMF system are presented in a block diagram. The common components of the setup are described in A. The audio tone generator mapped the target activity (control-point speed in CLMF) to graded audio tone frequencies. It generated audio signals at 44.1 kHz sampling based on the specified frequency and sent the signal to the audio output. The CLMF core was the primary program responsible for operating the CLMF system. It utilized config.ini, the camera factory, and DeepLabCut-Live(Kane et al. 2020), integrating the audio tone generator and reward delivery functions. This module also saved the data and configuration parameters with unique identifiers.

## Results

Consistent with prior studies demonstrating that mice can exert volitional control over their brain activity through feedback-based paradigms (Clancy et al. 2014; Luo et al. 2024; Neely, Koralek, et al. 2018; Clancy and Mrsic-Flogel 2021), our experiments show that mice learned to robustly modulate cortical signals when provided with closed-loop neurofeedback. Across training sessions, animals not only acquired the task (Figure 4A, initial rule RM ANOVA p=2.83e-5; Figure 4B, No-rule-change group RM-ANOVA p=9.6e-7) but also exhibited adaptability to altered feedback rules (Figure 4A, performance drop after rule change ANOVA, p=8.7e-9, new rule RM ANOVA p=8.3e-10), highlighting the flexibility of their learning. While feedback-driven brain control is well established, our work extends these findings by including multiple cortical regions, and by demonstrating that feedback based on movement-related signals can also drive learning despite the additional challenge of real-time behavioral tracking.

In our closed-loop experiment, mice were trained to associate a specific body movement with a reward. Over the course of 10 training sessions, the mice progressively learned the task, as evidenced by increased task performance across sessions (Figure 4A). The acquisition of this learned behavior was consistent with previous findings on motor learning, where reinforcement and task repetition lead to the refinement of motor skills (Wolpert, Diedrichsen, and Flanagan 2011; Krakauer et al. 2019). Importantly, the reward was provided 1 s after a successful trial to study the timing and nature of cortical activation in response to both the task and the reward. Concurrent with behavioral training, widefield cortical activity was recorded observing distinct patterns of neural activation that evolved as the mice learned the task.

We implemented and tested a compact, cross-platform system designed for closed-loop feedback experiments. Using this system, head-fixed mice successfully learned to associate either cortical activity or movement-derived signals with auditory feedback and water rewards. Performance increased steadily over sessions: in both one-ROI and two-ROI paradigms, animals showed a significant rise in the number of rewards obtained compared to their baseline (day 0) and control groups without feedback (Animations 1A–F). These results indicate that the system effectively links cortical dynamics or movement signals to behaviorally meaningful outcomes. Notably, both neurofeedback (CLNF) and movement-feedback (CLMF) modes produced reliable and graded closed-loop feedback.

In CLNF experiments, auditory tones mapped to ΔF/F_0_ values from cortical regions of interest accurately reflected ongoing neural activity (Figure 1). In two-ROI paradigms, differences between ROI activity were similarly mapped to sound frequency, and the commanded audio outputs were verified to be accurate across the full frequency range (Supplementary Figure 2). Improvements to the audio generation pipeline eliminated previous nonlinearities above 10 kHz, confirming that the feedback signal matched the intended output. Importantly, feedback latency was tested and found to be well within the range necessary for effective closed-loop learning. Using LED-triggering experiments (Supplementary Figure 1), we confirmed that the Raspberry Pi–based CLNF system delivered graded auditory feedback with an average latency of ∼63 ± 15 ms from event detection to output, using GPIO pins. Please note this latency calculation used GPIO pins for delay measurements, and there could be additional minimal overheads to generate the actual audio feedback signals. These findings demonstrate that our system provides rapid and accurate feedback sufficient to drive volitional control of brain or movement signals in head-fixed mice.

For the CLMF experiments, an Omron Sentech STC-MCCM401U3V USB3 Vision camera, connected to the Nvidia Jetson via its Python software developer kit (SDK), was used to calculate the feedback delay. The incoming stream of frames was processed in real-time using a custom deep-neural-network model that was trained using DeepLabCut (A. Mathis et al. 2018) and DeepLabCut-Live (Kane et al. 2020), designed to track previously defined points on the mouse. The model was incrementally improved by fine-tuning and re-training 26 times, using 2080 manually labeled frames spanning 52 videos of 10 mice. The pre-trained model, along with the data, is available at the link shared in the data and code availability statement in methods for anyone to use and fine-tune to adapt for similar platforms. The model was integrated into our CLMF program and deployed on an Nvidia Jetson device for real-time inference of tracked points. A Python-based virtual environment using Conda was created to install all software dependencies. The coordinates of the tracked points for each frame were appended to a list, forming a temporal sequence referred to as “tracks.” Upon offline analysis, these real-time tracks were found to be both accurate and stable throughout the duration of the videos.

To calculate the feedback delay for movements, a red LED was placed within the behavior camera’s field of view. Whenever a threshold-crossing movement was detected in the real-time tracking, the system triggered the LED to turn on. Temporal traces of the tracked left forelimb and the pixel brightness of the red LED were then extracted from the recorded video. By comparing these traces, the average delay between the detected movements and the LED illumination was calculated to be 67±20 ms, which represents the delay in the closed-loop feedback for the CLMF experiments.

For CLMF, mice were headfixed in a specially designed head-fixing chamber (Figure 1Biii, design guide and 3D model in methods) to achieve multiview behavioral recording from a single camera using mirrors (see methods section). In brief, mice are headfixed in a transparent rectangular tunnel (top view) with a mirror at the front (front view) and at the bottom (bottom view) that allows multiple views of the mouse body. The body parts that we track are: snout-top (snout in the top view), tail-top (base of the tail in the top view), snout-front (snout in the front view), FLL-front (left forelimb in the front view), FLR-front (right forelimb in the front view), snout-bottom (snout in the bottom view), FLL-bottom (left forelimb in the bottom view), FLR-bottom (right forelimb in the bottom view), HLL-bottom (left hindlimb in the bottom view), HLR-bottom (right hindlimb in the bottom view), tail-bottom (base of the tail in the bottom view) (Figure 1B). The rationale for selecting these body parts in a particular view was that they needed to be visible at all times to avoid misclassification in real-time tracking. By combining the tracks of a body part in different views, we can form a 3D track of the body part. In a 3D coordinate space having X, Y, and Z axes, tracked points (xb, yb) in the bottom view were treated as being in the 3D X-Y plane, and tracked points (xf, yf) in the front view were treated as being in the 3D Y-Z plane. Thus, X = xb, Y = yb, Z = yf formed tracked points in 3D for a given body part that was tracked in multiple views.

For example, FLL-front and FLL-bottom were tracking the left forelimb in front and bottom views, and by combining the tracks of these two points, we obtained a 3D track of the left forelimb (Figure 3E). Although these 3D tracks are available in real-time, for our CLMF experiments, we used 2D tracks (xb, yb) for behavioral feedback to keep the task simpler and comparable across mice. Audio output channels and GPIO pins on the Nvidia Jetson were used for audio feedback and reward delivery, respectively. Tracks of each body part were used to calculate the speed of those points in real-time, and a selected body part (also referred to as a control point) was mapped to a function generating proportional audio frequency (same as in CLNF, details in the method section). In the software we have developed, one can also choose to calculate acceleration, radial distance, angular velocity, etc., from these tracks and map it to the function generating varying audio frequency feedback. For our work, a range of spontaneous speeds was calculated from a baseline recording before the start of the training. The threshold speed to receive a reward was also calculated from the baseline recording and set at a value that would have yielded a reward rate of one reward per two-minute period (translates to a basal performance of 25% in our trial task structure). This thresholding step was done to allow the mice to discover task rules and keep them motivated.

**Figure 3.**
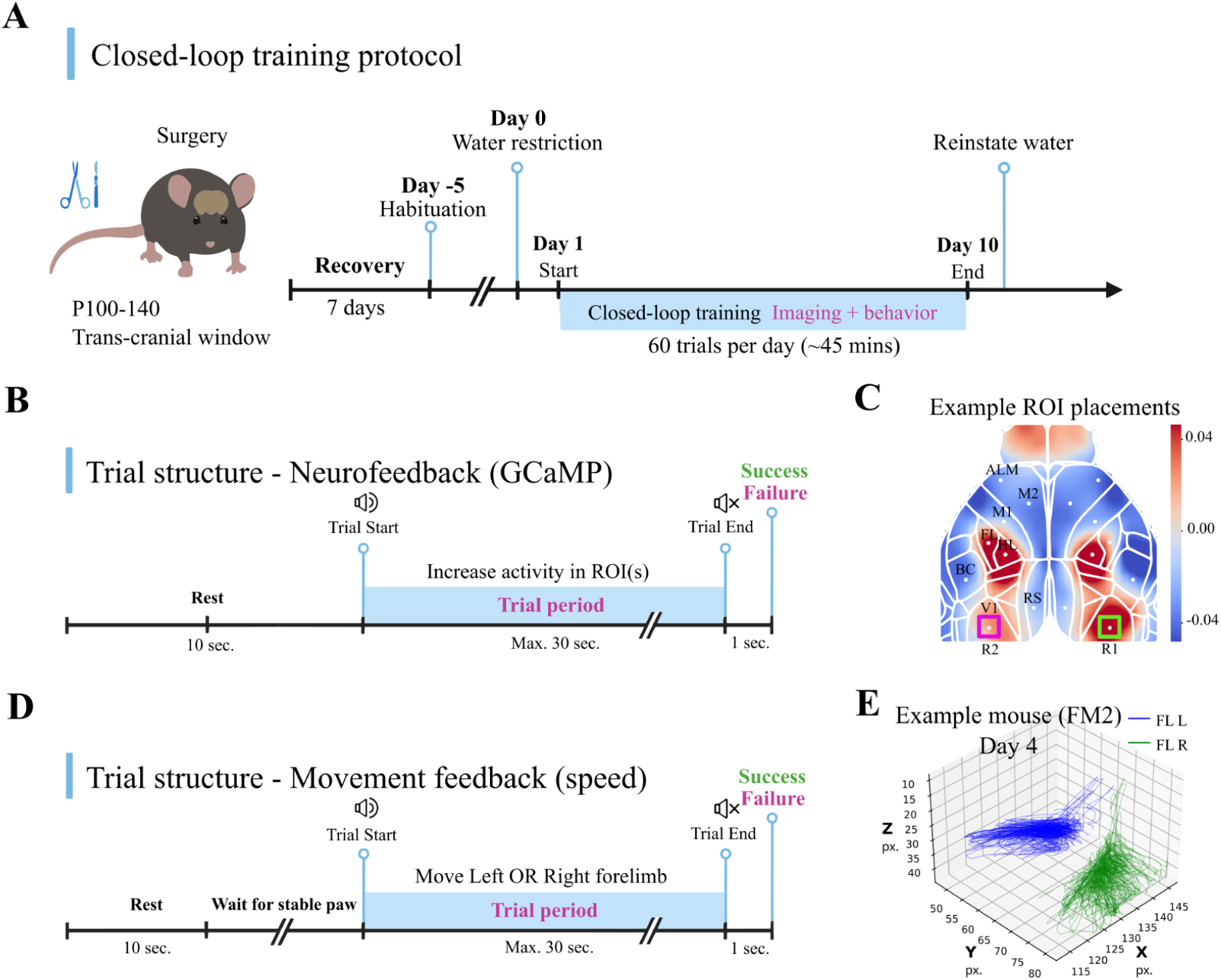
Experimental protocol and trial structure. A) Experimental Protocol (detailed in Methods): In brief, 90-day-old transgenic male and female mice were implanted with a transcranial window and allowed to recover for a minimum of 7 days. One day before the start of the experiment, the mice were placed on a water-restriction protocol (as detailed in Methods). Closed-loop experiment training commenced on day 1, during which mice were required to modulate either their target brain activity (GCaMP signals in regions of interest) or target behavior (tracked paw-speed) during daily sessions of approximately 45 minutes over the course of 10 days. Throughout this period, both cortical and behavioral activities were recorded. After the final experimental session, the mice were removed from the water-restriction protocol. B) Trial Structure of Cortical GCaMP-based Feedback Sessions: Each trial was preceded by a minimum of 10 seconds of rest, which was extended if the tracked body parts of the mouse were not stable (sum of changes across body parts > 1.5 mm i.e. 5 pixel values). Once the mouse remained stable and refrained from moving its limbs, the trial began with a basal audio tone of 1 kHz. The mice then had 30 seconds to increase rule-based activations (in the selected ROI) up to a threshold value to receive a water reward. A trial ended as soon as the activation reached the threshold, triggering a reward delivery for success, or timed out after 30 seconds with an overhead buzzer serving as a negative signal of failure. Both the reward and the negative signal were delivered within 1 second after the audio ceased at the end of each trial. C) Example dorsal cortical ΔF/F_0_ activity was recorded and overlaid with a subset of Allen CCF coordinates, which could be selected as the center of candidate regions of interest (ROIs) shown as Green and Pink color squares. D) Trial Structure of Behavior Feedback Sessions: The behavioral feedback trials followed a similar structure to the cortical feedback trials, with each trial preceded by at least 10 seconds of rest. The trial began with a basal tone of 1 kHz, which increased in frequency as the mouse’s paw speed increased. E) Example forelimb Tracking during Feedback Sessions: Forelimb tracking was performed in both the left (blue) and right (green) forelimbs using a camera coordinate system on day 4 of training with mouse FM2, which received feedback based on left forelimb speed. The forelimbs were tracked in 3D, leveraging multiple camera views captured using mirrors positioned at the bottom and front of the setup (Figure 1B).

Both CLNF and CLMF experiments shared a similar experimental protocol (surgery, habituation, then several days of training, Figure 3A). A daily session starts with a 30 sec rest period (no rewards or task feedback) followed by 60 trials (maximum 30 sec each) with a 10 sec rest between trials in CLNF, and a minimum of 10 sec rest or until tracked points are stable for 5 sec in CLMF (Figure 3B, 3D). After the habituation period, a spontaneous session of 30 minutes was recorded where mice were not given any feedback or rewards.

The spontaneous session was used to establish baseline levels of GCaMP activity (in target ROI(s) for CLNF experiments) or speed of a target body part for CLMF experiments. This was done to calculate the animal-specific threshold for future training sessions. A success in the trial resulted in a water drop reward that was delivered 1 sec after the end of the trial, and a failed trial ended with a buzzer vibrator 1 sec after the end of the trial (Figure 3B, 3D).

### Mice can explore and learn an arbitrary task, rule, and target conditions

CLNF training (Figure 1A) required real-time image processing, feedback, and reward generation. Feedback was a graded auditory tone mapped to the relative changes in selected cortical ROI(s) or movement speed of a tracked body part. Training was conducted using multiple sets of cortical ROI(s) (list of ROI rules and their description can be found in Table 3 and visualized in Supplementary Figure 7) on both male and female mice (see Table 5 for list of mice and their task rules), wherein the task was to increase the activity in the selected ROI(s), according to the rule (also referred to as ‘the rule’ in the future), to exceed a predetermined threshold (Figure 6A, 6B). Fluorescence activity changes (ΔF/F_0_) were calculated using a running baseline of 5 seconds, with single or dual ROIs on the dorsal cortical map selected as targets. The distribution of target ROI(s) activity on day 9 of training was higher than on day 1 (Figure 5A). A linear regression analysis of whole-session ΔF/F_0_ activity in the 2-ROI experiment revealed a shift in the regression line slope, favoring the ROI that was required to increase according to the task rule (Figure 6C). In general, all ROIs assessed (Table 3, Supplementary Figure 7) that encompassed sensory, pre-motor, and motor areas were capable of supporting increased reward rates over time based on analysis of group and pooled ROI data (Figure 4A, Animation 1). A ΔF/F_0_ threshold value was calculated from a baseline session on day 0 that would have allowed 25% performance. Starting from this basal performance of around 25% on day 1, mice (CLNF No-rule-change, N=23, n=60 and CLNF Rule-change, N=17, n=60) were able to discover the task rule and perform above 80% over ten days of training (Figure 4A, RM ANOVA p=2.83e-5), and Rule-change mice even learned a change in ROIs or rule reversal (Figure 4A, RM ANOVA p=8.3e-10, Table 5 for different rule changes). There were no significant differences between male and female mice (Supplementary Figure 3A). To investigate whether certain ROI(s) were better than others in terms of performance, we performed linear regression of the success rate over the days and, based on the slope of the fitted line, discovered ROI rules that yielded statistically different progression (fast and relatively slower) of success rate from the mean slope of all ROIs (Supplementary Figure 3C). We visualized these significantly different ROI-rule-based success rates and segregated them into fast and slow based on mean slope (>=0.095 slope was designated fast, else slow) of the progression (Supplementary Figure 3D). Our analysis revealed that certain ROI rules (see description in methods) lead to a greater increase in success rate over time than others (Supplementary Figure 3D). As mice learned the CLNF task and their performance improved, task latency progressively decreased (Figure 4C). Here, task latency refers to the time required by the mice to complete the task within the 30-s trial window. For this measure, we included all trials, both successful and unsuccessful. In the 2-ROI experiment where the task rule required, for example, “ROI1 - ROI2” activity to cross a threshold for reward delivery, mice displayed divergent strategies. Some animals predominantly increased ROI1 activity, whereas others reduced ROI2 activity, both approaches could lead to successful threshold crossing (Figure 11; see average reward centered responses).

**Figure 4:**
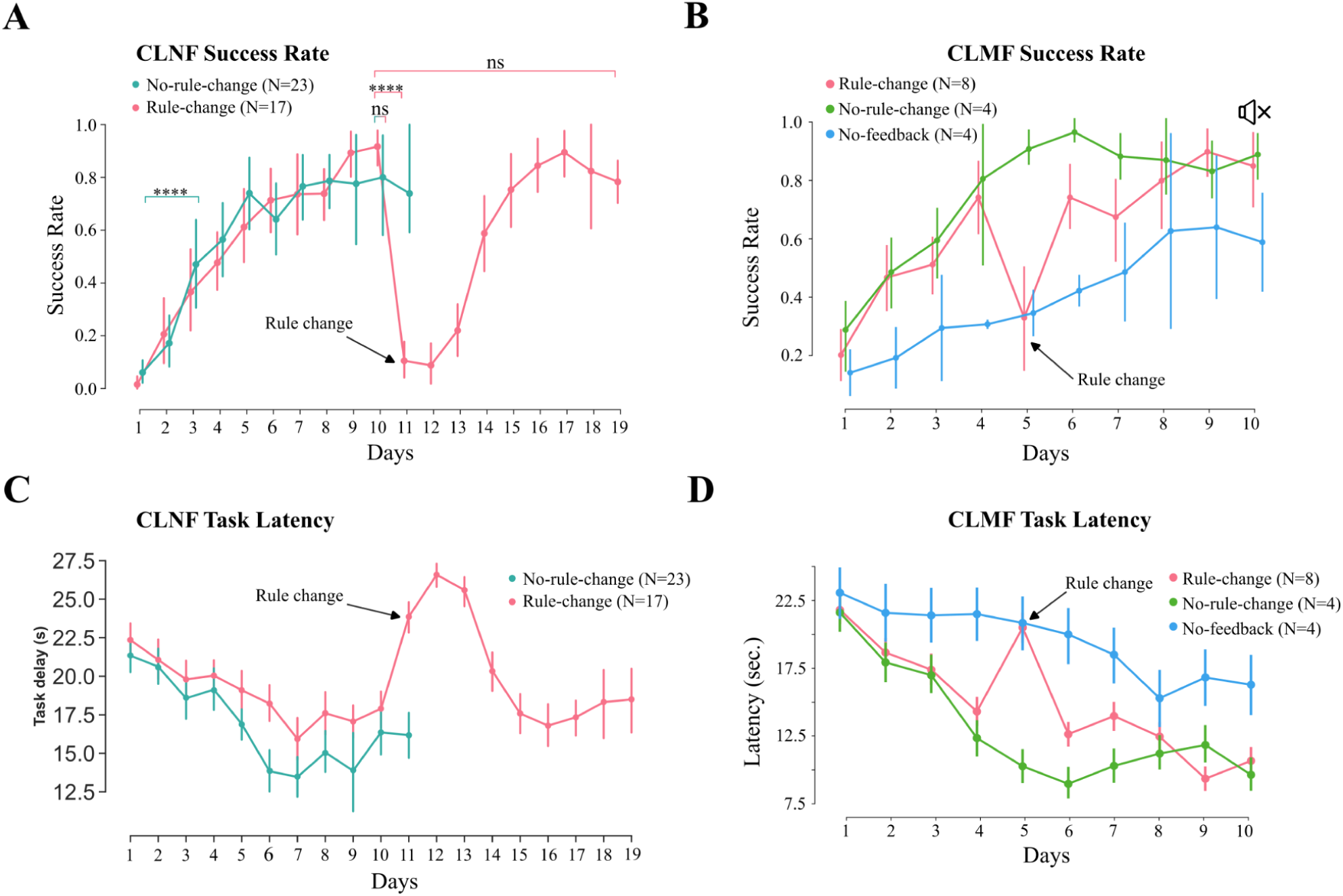
Closed-loop feedback helped mice learn the task and achieve superior performance in CLNF and CLMF in both experiments. A) No-rule-change (N=23, n=60, in Table 5) and Rule-change (N=17, n=60) mice were able to learn the CLNF task over several sessions, with performance above 70% by the 10th session (RM ANOVA p=2.83e-5). The rule change (in pink, day 11) led to a sharp decline in performance (ANOVA, p=8.7e-9), but the mice were able to adapt and learn the task rule change (RM ANOVA p=8.3e-10; see Table 5 for different rule changes). The method to determine the ROI(s) used in the changed task rule is described in the methods section. B) Three groups were employed for CLMF experiments. The “Rule-change” group (N=8, n=60 received feedback, in pink) was trained with task rule mapping auditory feedback to the speed of the left forelimb and was able to perform above a 70% success rate in four days. The task rule mapping was changed from the left to the right forelimb on day 5, so the rewards as well as audio frequencies would now be controlled by the right forelimb. The “No-rule-change” group (N=4, n=60 received audio feedback, no rule change, in green) and the “No-feedback” group (N=4, n=60 no graded audio feedback, no rule change, in blue) were control groups to investigate the role of audio feedback. The performance of the “No-feedback” mice, who did not receive the graded feedback, was never on par (RM-ANOVA p=0.49) with the “No-rule-change” group that received the feedback (RM-ANOVA p=9.6e-7). Additionally, to test the effect of auditory feedback on mice that already had learned the task, we turned off the auditory feedback on day 10 for all mice (indicated by the speaker with a cross). There was no significant change in performance due to feedback removal, indicating that feedback was not necessary once the task was learned. C) Task latencies in each group follow the trend of their performance. Rule change (N=8) and no rule change (N=4) task latencies gradually came down, with an exception on day 5 for rule change when the task rule was changed. No feedback (N=4) task latencies are never on par with the groups that received feedback. D) CLMF Rule-change (N=8) behavior, we looked at the maximum speeds of the left and right forelimbs. The paw with the maximum speed follows the task rule and switches with the change in the task rule. It is worth noting that the task was not restricted to other body parts; i.e., the mice were free to move other body parts along with the control point.

**Figure 5:**
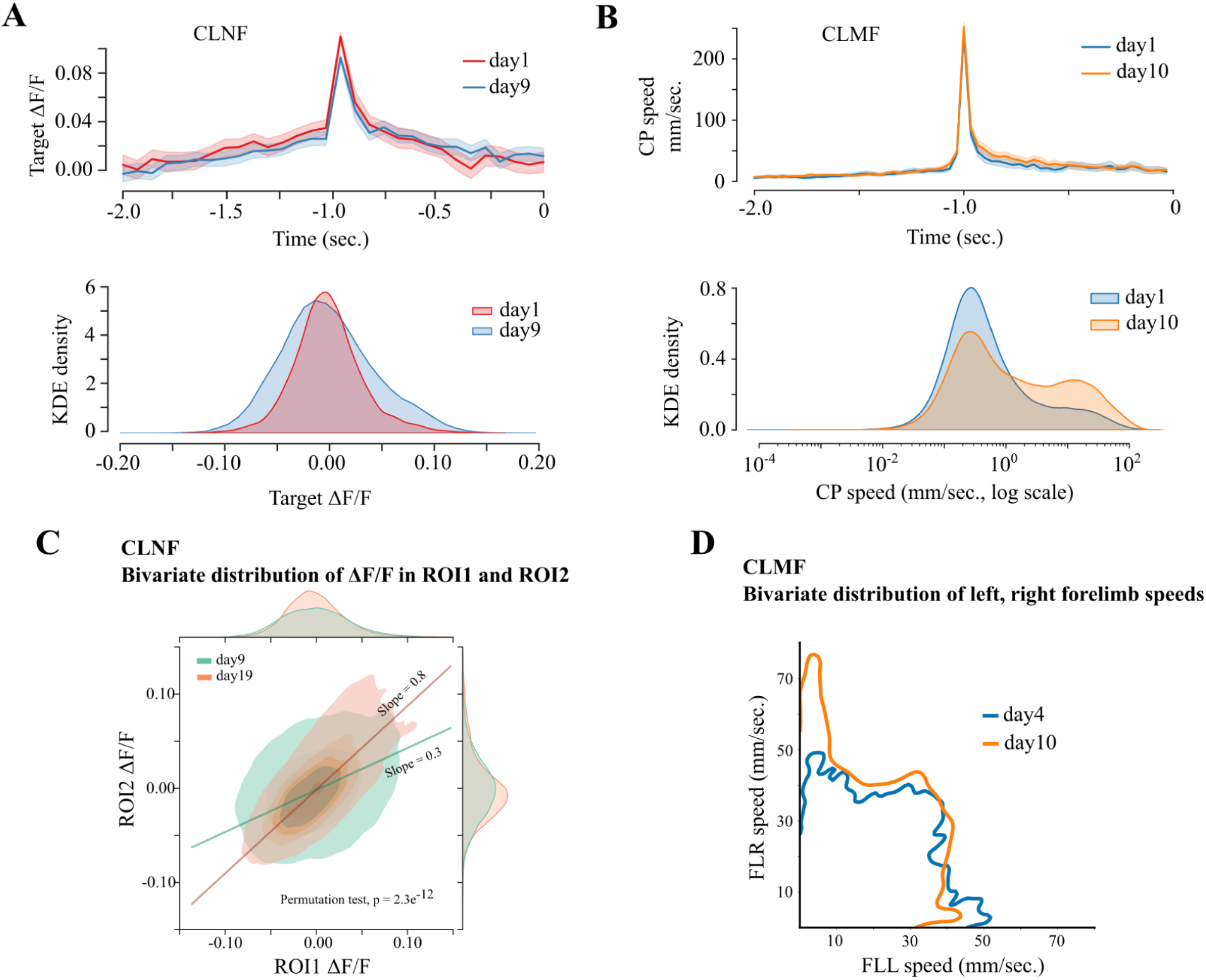
Closed-loop feedback helped mice learn the task and achieve superior performance in CLNF and CLMF in both experiments. A) Reward-aligned average (N=4) ΔF/F_0_ signals associated with the target rule on day1 and day9 (top plot). Kernel density estimate (KDE) of target ΔF/F_0_ values during the whole session on day1 and day9 of 1-ROI experiments (bottom plot). B) Reward-aligned average (N=4) target paw speed on day1 and day10 (top plot). Kernel density estimate (KDE) of target paw speeds on day1 and day10 (bottom plot). C) In the context of CLNF 2-ROI experiments, bivariate distribution of ROI1 and ROI2 ΔF/F_0_ values during whole sessions on day9 and day19, with densities projected on the marginal axes. The task rule on day9 was “ROI1-ROI2 > thresh.” as opposed to “ROI2-ROI1 > thresh.” on day19. The bivariate distribution is significantly different (Multivariate two-sample permutation test, p=2.3e-12) on these days, indicating a robust change in activity within these brain regions. D) Joint (bivariate) distribution of left and right paw speeds during the whole session on day4 and day10 of CLMF. Left and right forelimbs were control-point (CP) on day4 and day10 respectively. There is a visible bias in the bivariate distribution towards the CP on respective days.

**Figure 6:**
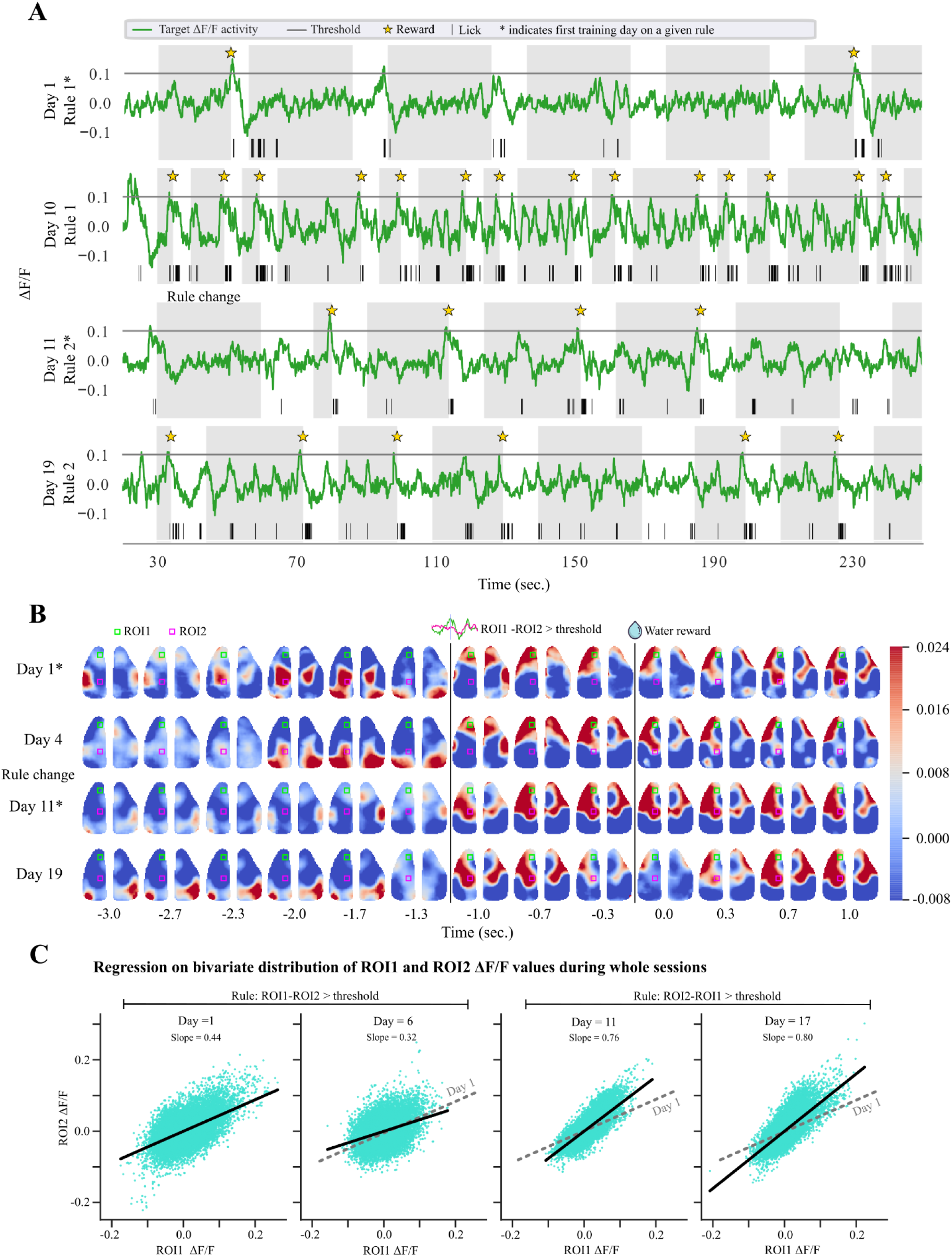
CLNF: Cortical activity during the closed-loop-neurofeedback training. A) Target-rule-based ΔF/F_0_ traces in green on day 1 with rule-1 (top row), on day 10 with rule-1 (second row), on day 11 with new rule-2 (third row), and on day 19 with rule-2 (fourth row). Shared regions are trial periods and regions between grey areas are rest periods. The grey horizontal line depicts the threshold above which mice would receive a reward. Golden stars show the rewards received, and short vertical lines in black show the spout licks. B) Representative reward-centered average cortical responses of the 2ROI experiment on labeled days. ROI1 (green) and ROI2 (pink) are overlaid on the brain maps. The task rule on day 1 and day 4 was “ROI1-ROI2 > thresh,” as opposed to “ROI2-ROI1 > thresh” on day 11 and day 19. C) Linear regression on ROI1 and ROI2 ΔF/F_0_ during whole sessions for an example mouse. The regression fit inclines towards ROI1 in sessions where the rule was “ROI1-ROI2 > thresh.” (day1 slope=0.44, day6 slope=0.32) while it leans towards ROI2 after the task rule switches to “ROI2-ROI1 > thresh.” (day11 slope=0.76, day17 slope=0.80).

**Figure 7:**
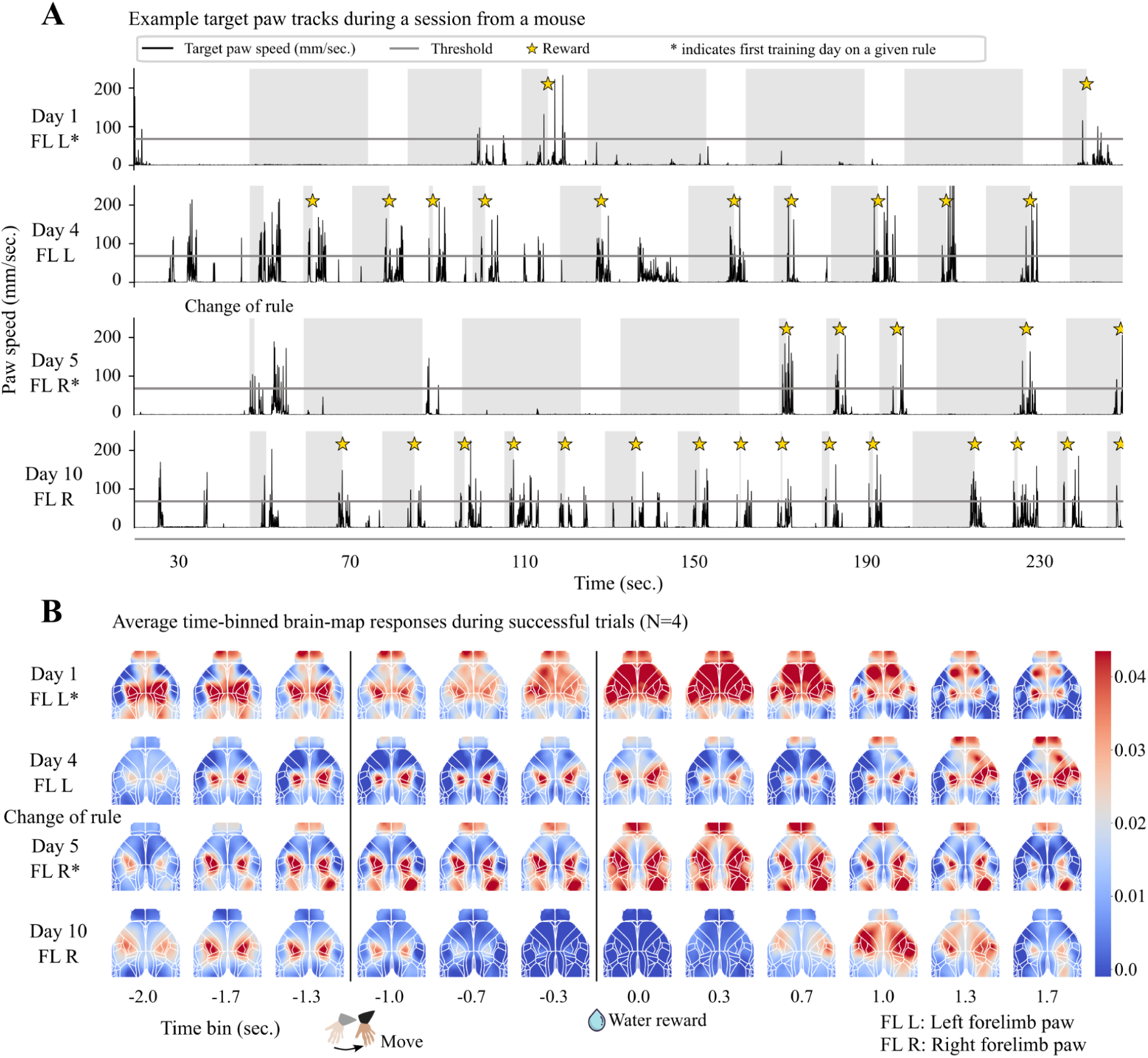
CLMF: Speed of the tracked target body part and cortical activity. A) Left forelimb speed (black), target threshold (grey line), and rewards (golden stars) during a sample period in a session on day 1 of the closed-loop training (top row). Shaded regions are trial periods with interspersed rest periods in white. Left forelimb speed, and rewards on day4 (second row). The target body part was changed from the left forelimb to the right forelimb on day5 (third row). Thus, day5 is the first training day with the new rule. Right forelimb speed, and rewards on day 10 of the training (fourth row). B) Reward-centered average cortical responses on days corresponding to rows in A. The target threshold was crossed at -1 s, and the reward was delivered at 0 s. Notice the task rule change on day 5.

**Table 1:**
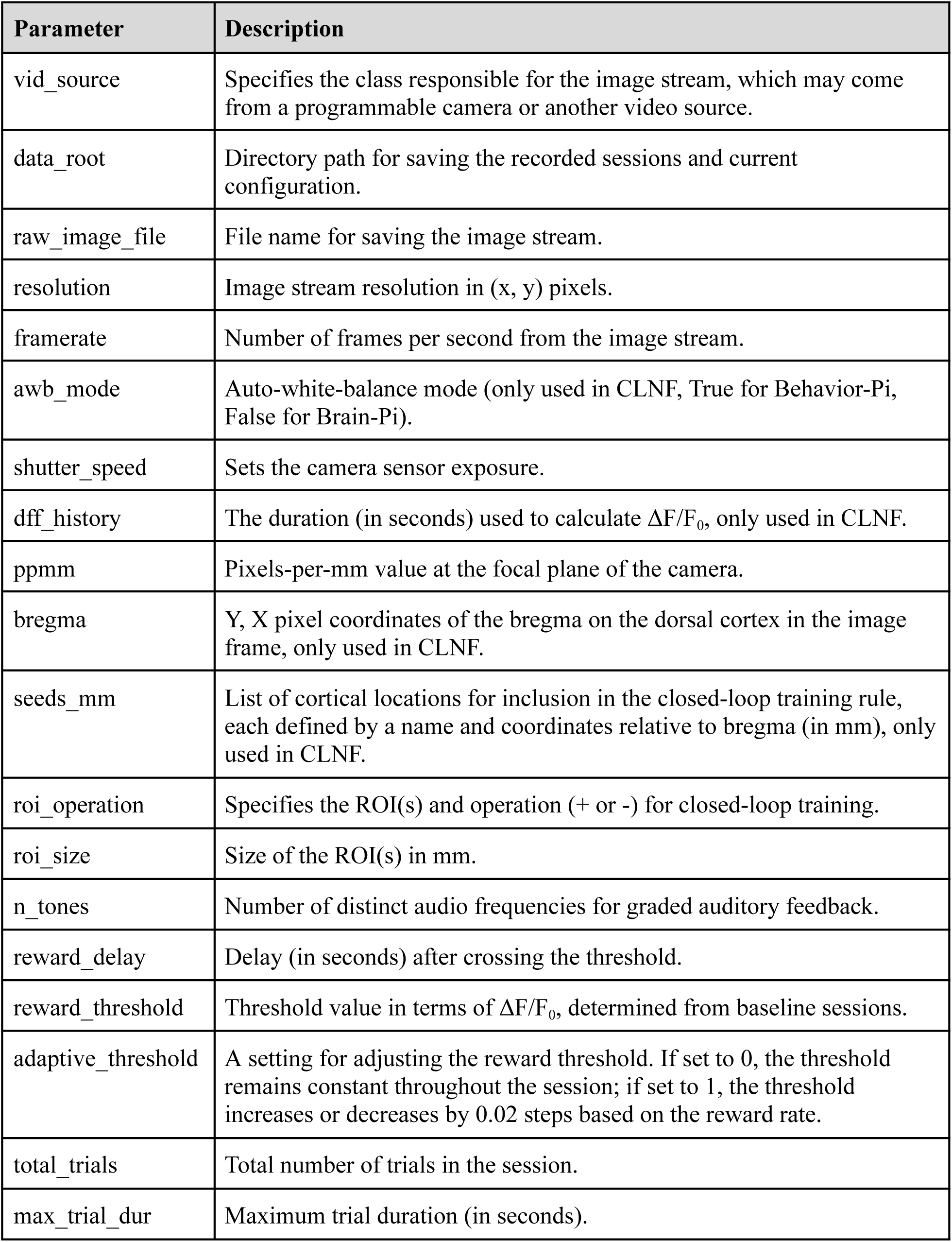
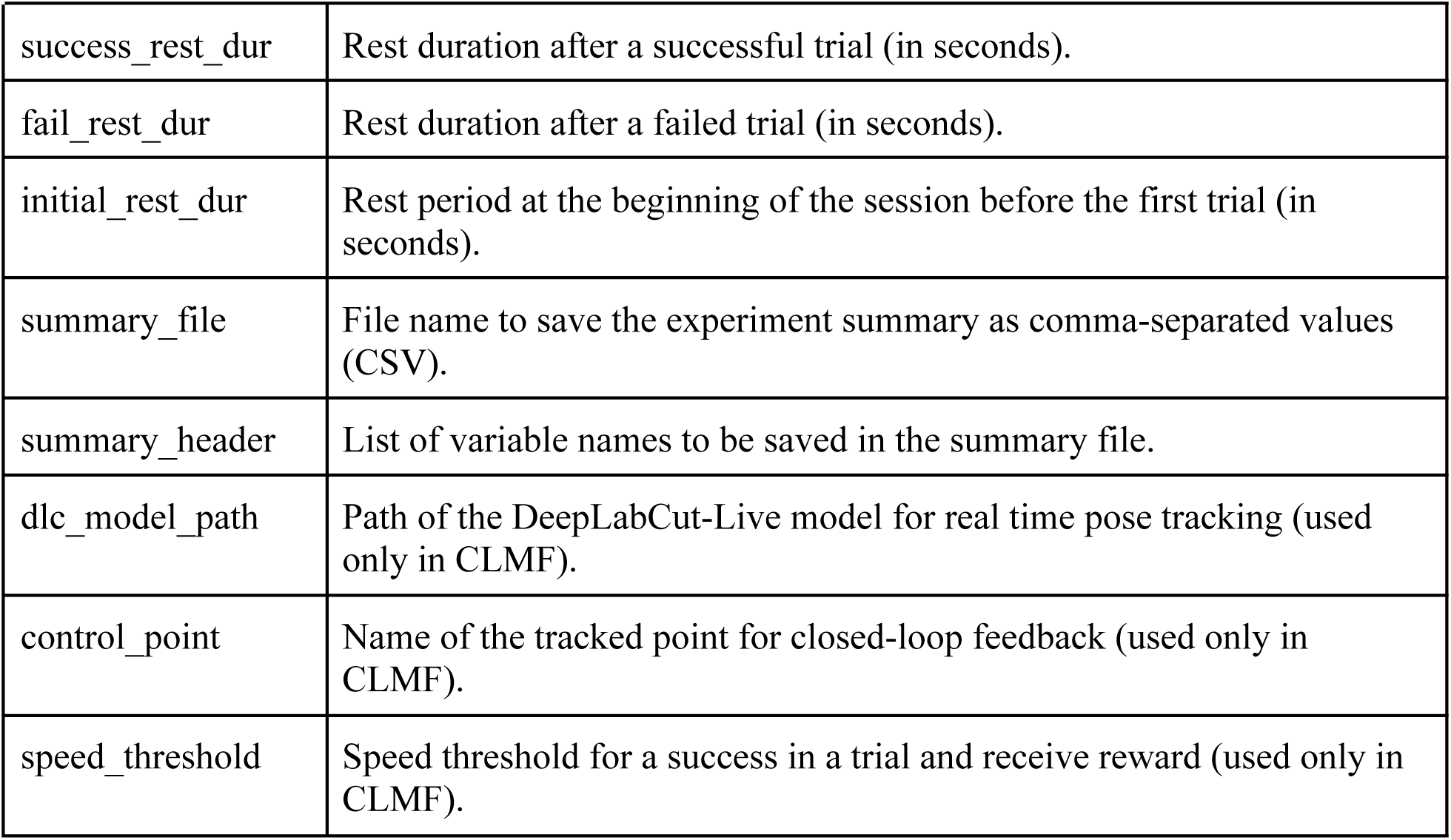
Key configuration parameters in CLoPy.

In CLMF training, the real-time speed of a selected (also referred to as control point (CP) in terms of tracked points) point was mapped to the graded audio tone generator function. We trained the mice with the FLL bottom as the CP for auditory feedback and reward generation (example trial in Animation 2). Mice reached 80% performance (Figure 4B, CLMF Rule-change, No-rule-change) in the task within four days of training (RM ANOVA, p = 8.03e-7). They were eliciting the target behavior (i.e., moving CP at high speed) more frequently on later days compared to the first day of training (Figure 5B), and the correlations of CP speed profiles became more pronounced during the trial periods as compared to the rest period (Figure 8A).

**Figure 8:**
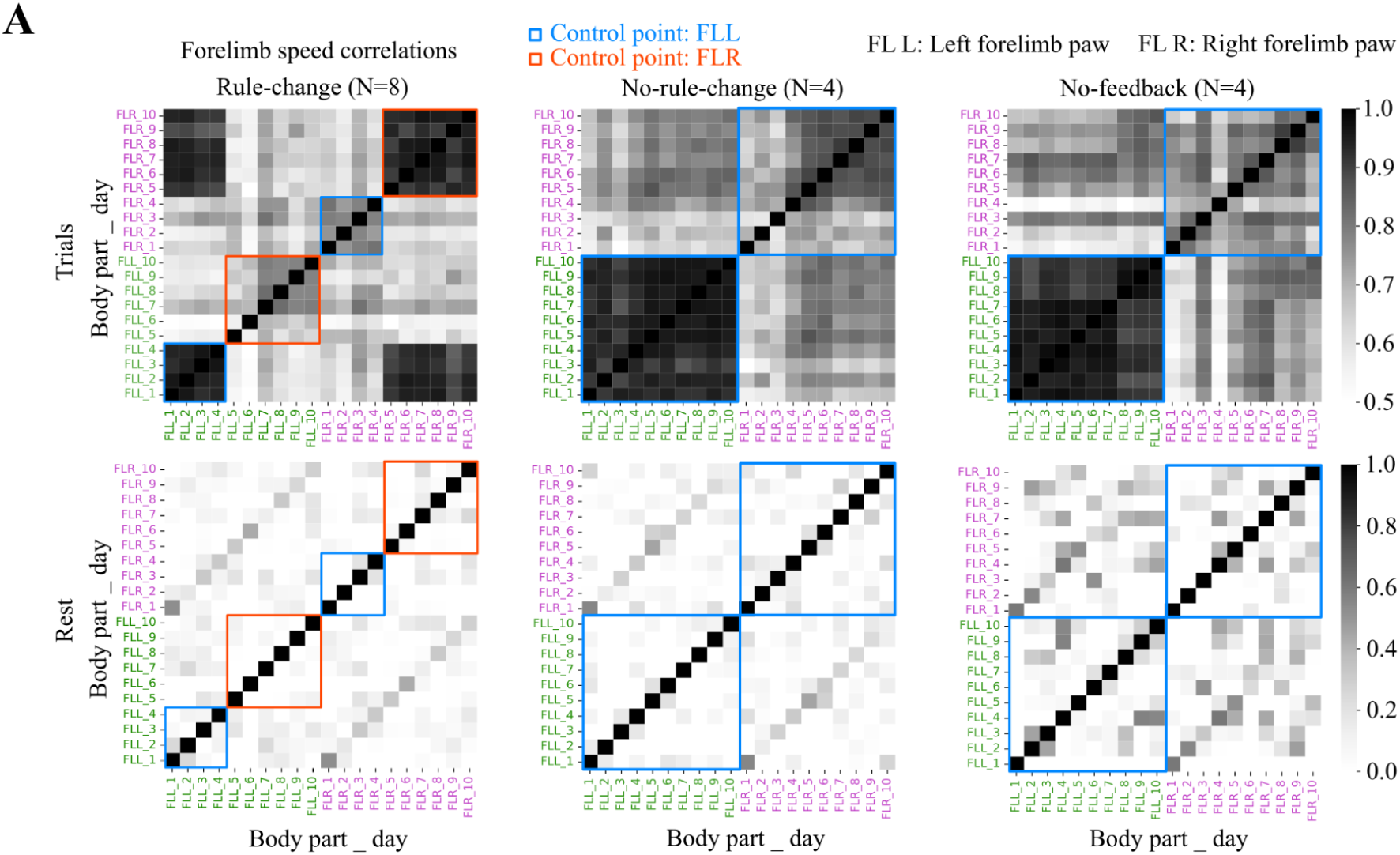
CLMF: Correlation matrix showing pairwise correlations of left and right forelimb speed profiles. A) Top row: During rewarded trials, over the training sessions of CLMF Rule-change (left), No-rule-change (middle), and No-feedback (right). High correlations (dark cells) between speed profiles of CP indicate a unilateral bias in the movement. It is worth noting the drastic changes in correlations as the control-point was changed from left forelimb to right forelimb in Rule-change mice on day4. Bottom row: During rest periods, over the training sessions of CLMF Rule-change (left), No-rule-change (middle), and No-feedback (right).

### Mice can rapidly adapt to changes in the task rule

We have assessed how mice respond to changes in closed-loop feedback rules. CLNF Rule-change mice went through a change in regulated cortical ROI(s) (Figure 6A, 6B) after being trained on an initial set of ROI(s) (Table 5, Rule-change). The new ROI(s) were chosen from a list of candidate ROI(s) for which the cortical activations were still low (see “Determining ROI(s) for change in CLNF task rule” under Methods). Reward threshold values were then recalculated for the ROI(s). A full list of ROI(s) we tried in separate mice is listed in Table 5. Mouse performance degraded on the first day of the rule switch to the new ROIs (ANOVA, p=8.7e-9), compared to the previous day (Figure 4A), and quickly recovered within five days (RM ANOVA p=8.3e-10) of further training. All new cortical ROI(s) appeared to support recovery to the pre-perturbation success rate (Table 5), and data were pooled across mice. This can also be observed in the bivariate distribution of a whole session ΔF/F_0_ values in ROI1 vs ROI2 on day 9 (before rule change) and day 19 (last day after rule change) in Figure 5C. Additionally, the linear regression analysis of whole-session ΔF/F_0_ activity after rule change in the 2-ROI experiment revealed a shift in the regression line slope, favoring the ROI that was required to increase according to the new task rule (Figure 6C). Following the rule change, CLNF task latency increased substantially and persisted at a high level across subsequent sessions, eventually declining as task performance adapted to the new rule (Figure 4C).

Similarly, in the CLMF experiments, the target body part was changed from FLL to FLR for Rule-change mice on day 5 and was associated with a significant drop in their success rate from 75-80% to ∼40% (ANOVA p=0.008, Figure 4B). Surprisingly, mice quickly adapted to the rule change and started performing above 70% on day 6 (example trial in Animation 3). Looking closely at their reward rate on day 5 (day of rule change), they had a higher reward rate in the second half of the session as compared to the first half (Supplementary Figure 4A), indicating they were adapting to the rule change within one session. This can also be observed in the bivariate distribution of left and right paw speeds on day 4 (before rule change) and day 10 (last day after rule change) in Figure 5D. As the mice were learning the CLMF task (increasing performance), task latency decreased (Figure 4D). Task latency in this context is the time taken by mice to perform the task within a trial period of 30 s. We included all trials, both successful and unsuccessful, in our calculation. Given that the maximum trial duration is 30 s, the task latency was capped at 30 s. Following the trend in task performance, the task latency started decreasing during the first four days but increased on day 5 (rule change) and started to drop again afterward.

We also examined the average paw speeds and distributions during trial periods. It is worthwhile noting that for the rule-change group, left forelimb speeds were higher than right forelimb speeds from day 1 to day 4 (when task rule was to move left forelimb). When the rule was switched from left forelimb to right forelimb on day 5, left forelimb speeds dropped below the right forelimb speeds (Supplementary Figure 4E).

### Graded feedback helps in task exploration during learning, but not after learning

To investigate the role of audio feedback in our task, we also had a group with no task-related graded audio feedback (CLMF No-feedback) and instead received audio with a constant frequency (1 kHz) throughout the trials. CLMF No-rule-change mice who received continuous graded auditory feedback significantly improved their task performance (CLMF No-rule-change RM-ANOVA p = 9.6e-7) and outperformed the CLMF No-feedback mice (Figure 4B) very early (No-feedback RM-ANOVA p = 0.49), indicating the positive role of graded feedback for task exploration and learning the association. When graded audio feedback was removed for CLMF Rule-change mice on day 10, it did not affect their task performance, indicating the feedback was not essential to keep performing the task that they had already learned.

### Cortical responses became focal and more correlated as mice learned the rewarded behavior in CLMF

There are reports of cortical plasticity during motor learning tasks, both at cellular and mesoscopic scales (Makino et al. 2017; Huber et al. 2012; W. E. Allen et al. 2017). As mice become proficient in a task, their brain activity becomes more focused and less widespread (Figure 9). We noticed the peak activity in different regions of the cortex around the rewarding movement decreased ΔF/F_0_ in amplitude over days (Figure 9A, 9B, 9C, 9E). To quantify this, we measured the peak ΔF/F_0_ (ΔF/F_0_peak) value in the time window from -1s to +1s relative to the body part speed threshold crossing event in each cortical region. Consistent with our observations, ΔF/F_0_peak in several cortical regions gradually decreased over sessions, including the olfactory bulb, sensory forelimb, and primary visual cortex (Figure 9A, 9B, 9C). Notably, when the task rule was changed from FLL to FLR in the CLMF Rule-change mice, we observed a significant increase in ΔF/F_0_peak in regions including the olfactory bulb (OB), forelimb cortex (FL), hindlimb cortex (HL), and primary visual cortex (V1), as shown in Figure 9. We believe the decrease in ΔF/F_0_peak is unlikely to be driven by changes in movement, as movement amplitudes did not decrease significantly during these periods (Figure 9D CLMF Rule-change). However, the decrease in ΔF/F_0_peak followed the same trend as task latency (Figure 4D), suggesting that the decrease in ΔF/F_0_peak is especially prominent in trials where mice were prepared to make the fast movement.

**Figure 9:**
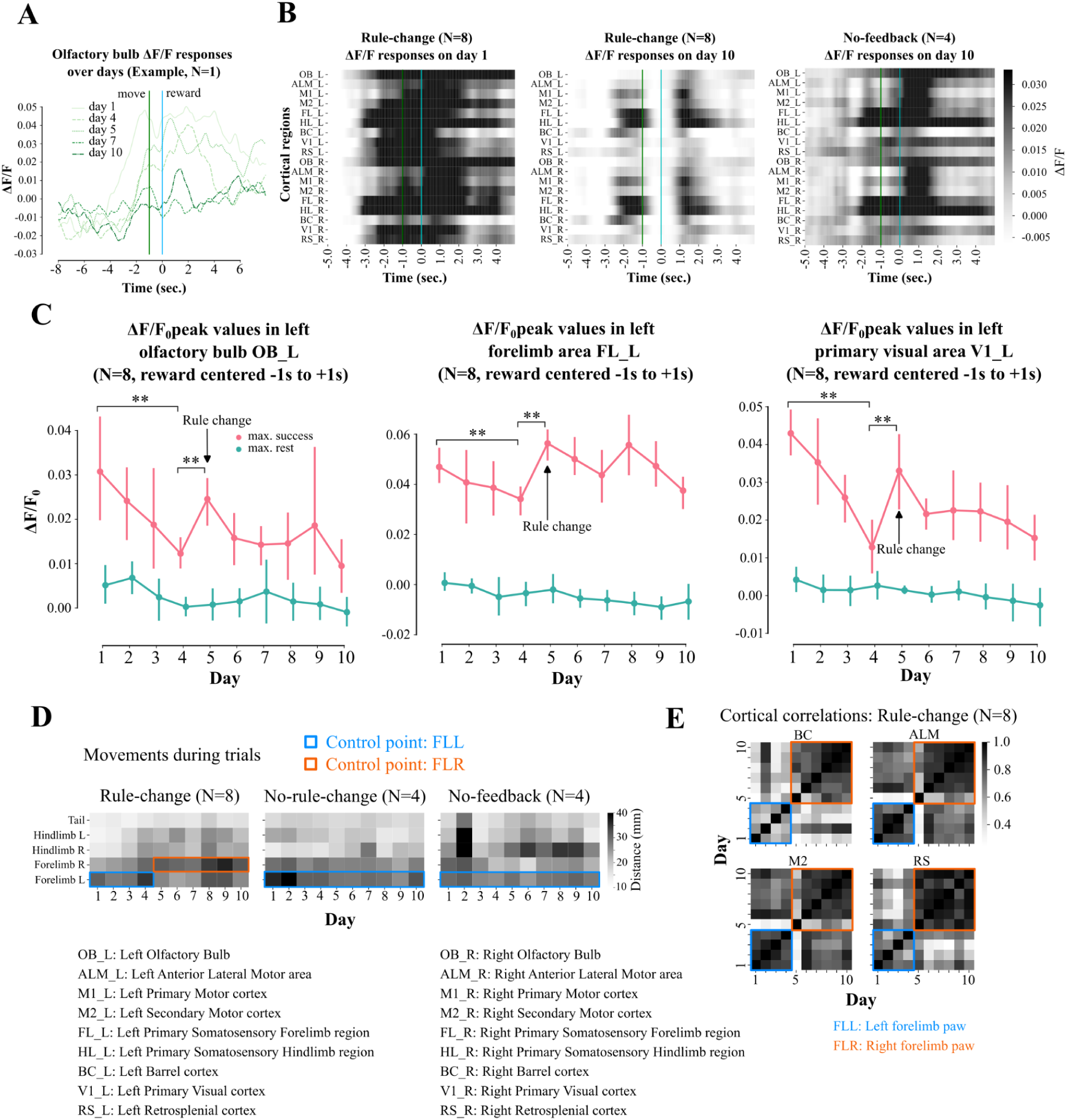
Cortical dynamics and network changes during longitudinal CLMF training. A) Reward-centered average responses in the olfactory bulb decrease over the days as performance increases. Data shown is a representative example from sessions of mouse. B) Cortical responses become focal and closely aligned to the paw movement (green line) and reward (cyan line) events on day 10 for group-1 (received feedback) as compared to group-3 (no-feedback). C) ΔF/F_0_ peak values during successful trials (pink) and during rest (cyan) over the ten-day training period in the olfactory bulb (OB) (left, day 1-day 4 p-value=0.025, day 4-day 5 p-value=0.008), forelimb area (FL) (center, day 1-day 4 p-value=0.008, day 4-day 5 p-value=0.04), and primary visual cortex (right, day 1-day 4 p-value=0.04, day 4-day 5 p-value=0.002). Statistical significance was assessed using the Mann-Whitney test and corrected for multiple comparisons with the Benjamini-Hochberg procedure. D) Average movement (mm) of different tracked body-parts during trials in CLMF Rule-change (left), No-rule-change (center), No-feedback (right). E) Correlations between cortical activation on each training session in barrel-cortex (BC, top left), anterolateral motor cortex (ALM, top right), secondary motor cortex (M2, bottom left), and retrosplenial cortex (RS, bottom right).

These results suggest that motor learning led to less cortical activation across multiple regions, which may reflect more efficient processing of movement-related activity. In addition to ΔF/F_0_peak reflecting signs of potentially more efficient cortical signaling, intracortical GCAMP transients measured over days became more correlated (to each other) during the trials as the task performance increased over the sessions (Figure 9E). Analysis of pairwise correlations between cortical regions (referred to as seed pixel correlation maps) revealed distinct network activations during rest and trial periods (Figure 10). While the general structure of the correlation maps remained consistent between trial and rest periods, certain correlations, such as between V1 and RS, were heightened during task trials, whereas others, such as between M1, M2, FL, and HL, consistently increased over the training sessions (Figure 10).

**Figure 10:**
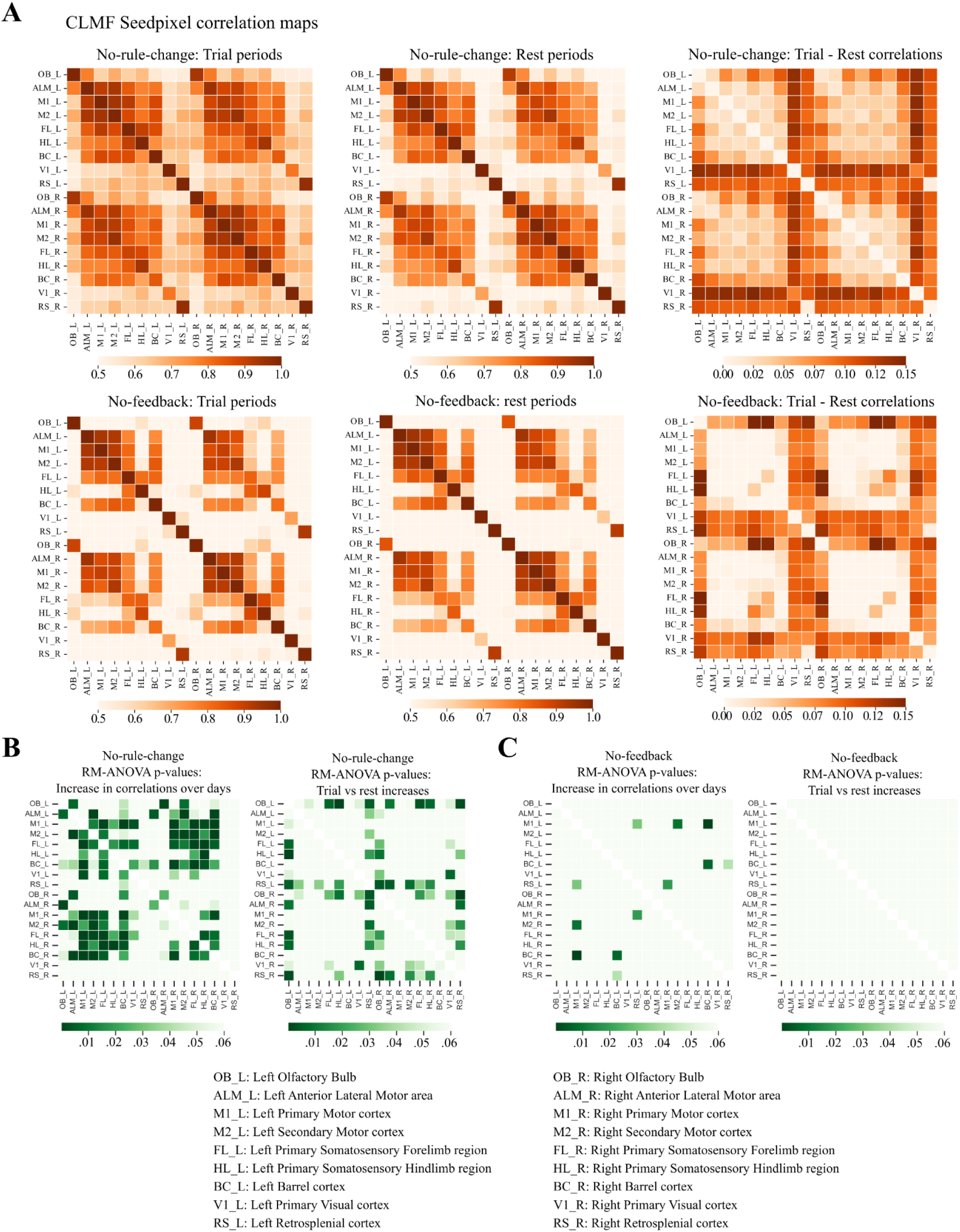
CLMF cortex-wide seed pixel correlation maps. Pairwise correlations between activity at cortical locations (also referred to as seed pixel locations). A) Top row: No-rule-change average seed pixel correlation map during trial periods (left), during rest periods (middle), and difference of average correlation map during trial and rest (right). Bottom row: No-feedback average seed pixel correlation map during trial periods (left), during rest periods (middle), and difference of average correlation map during trial and rest (right). B) Significant increase in pairwise seed pixel correlations as RM-ANOVA p-value (Bonferroni corrected) matrix between training sessions over the days (left) and between trial vs rest periods (right) for CLMF No-rule-change mice. C) Significant increase in pairwise seed pixel correlations as RM-ANOVA p-value (Bonferroni corrected) matrix between training sessions over the days (left) and between trial vs rest periods (right) for CLMF No-feedback mice.

**Figure 11:**
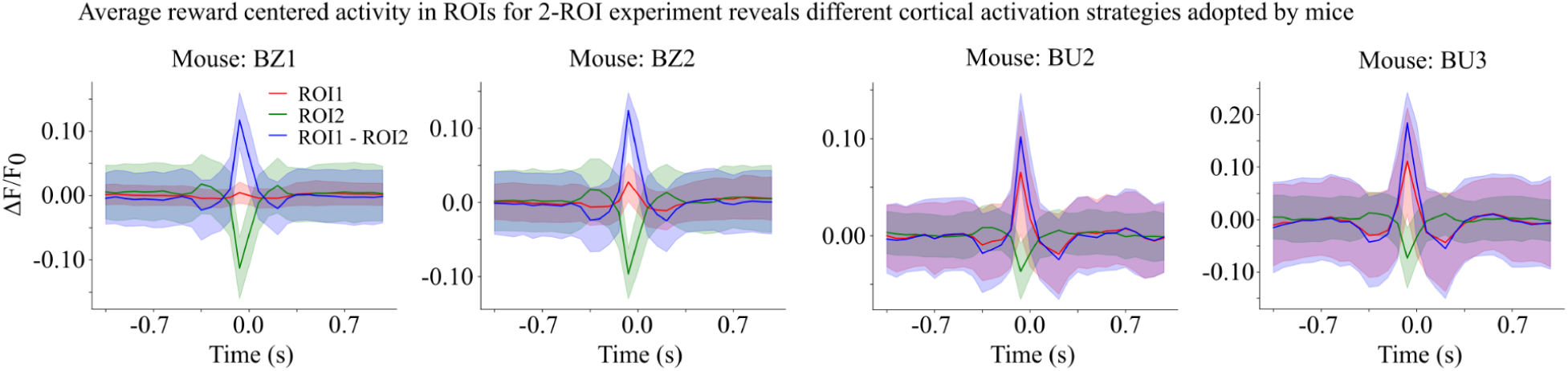
Average reward centered activity in ROIs for 2-ROI experiment. Examples of reward centered activity in both the ROIs in a 2 ROI experiment reveals different strategies adopted by mice. For a task rule specifying “ROI1 - ROI2 > threshold”, mouse BZ1 and BZ2 decreased ROI2 activity to meet the reward threshold while mouse BU2 and BU3 increased ROI1 activity.

To statistically examine the differences in seed pixel correlations during trials and rest periods, as well as how these correlations changed over training sessions (Day 1-10), we conducted a two-way repeated measures ANOVA (RM-ANOVA) on the seed pixel correlation maps for each day. The two variables for the RM-ANOVA were experimental condition (trial vs. rest) and session number (Day 1-10). This analysis generated two distinct matrices of Bonferroni-corrected p-values (Figure 10), one corresponding to each variable, which segregated the seed pixel correlations that were different between trial and rest periods and those that changed over the training sessions.

### Distinct task- and reward-related cortical dynamics

During the early sessions (days 1 to 3), cortical activity was observed to be spatially widespread and engaged multiple cortical regions. Temporally, the activity spanned both task-related and reward-related events, with no clear distinction between the two phases (Figure 9B, left). This broad activation pattern is consistent with the initial stages of learning, where the brain recruits extensive cortical networks to process novel tasks and integrate sensory feedback (Peters, Chen, and Komiyama 2014).

As the mouse performance improved in the later sessions (Days 8 to 10), the cortical activity became more segregated both spatially and temporally (Figure 9B, middle). This segregation was particularly notable in mice that received closed-loop feedback (Rule-change), where the spatiotemporal patterns of cortical activation were more closely aligned with the specific task and reward events. This transition from widespread to segregated activation is indicative of the brain’s optimization of neural resources as the task becomes more familiar, a phenomenon that has been reported in studies of skill acquisition and motor learning (Makino et al. 2017; Huber et al. 2012). Previous studies have shown that feedback, especially when provided in a temporally precise manner, can accelerate the cortical plasticity associated with learning (Ganguly and Carmena 2009). Overall, these findings highlight the importance of closed-loop feedback in motor learning and suggest that real-time neurofeedback can enhance the specificity of cortical representations, potentially leading to more efficient and robust learning outcomes.

## Discussion

### Flexible, cost-effective, and open-source system for a range of closed-loop experiments

We developed a versatile, cost-efficient, and open-source platform designed to support a wide array of closed-loop neuroscience experiments, including closed-loop neural feedback (CLNF) and closed-loop movement feedback (CLMF). This system enhances accessibility for the broader research community, enabling the execution of complex experimental paradigms with minimal resource constraints. Our system is built on readily available hardware components, such as Raspberry Pi and Nvidia Jetson platforms, and leverages Python-based software for real-time data processing, analysis, and feedback implementation. The modular approach ensures that the system can be easily customized to meet the specific requirements of different experimental paradigms. Our study demonstrates the effectiveness of a versatile and cost-effective closed-loop feedback system for modulating brain activity and behavior in head-fixed mice. By integrating real-time feedback based on cortical GCaMP imaging and behavior tracking, we provide strong evidence that such closed-loop systems can be instrumental in exploring the dynamic interplay between brain activity and behavior. The system’s ability to provide graded auditory feedback and rewards in response to specific neural and behavioral events showcases its potential for advancing research in neurofeedback and behavior modulation.

The hardware backbone of our system is designed around the Raspberry Pi 4B+ and Nvidia Jetson devices, chosen for their compactness, low cost, high computational power, and wide community support. These platforms have been demonstrated to be effective in neuroscience research, for several applications (Michelson et al. 2023; Murphy et al. 2020; Silasi et al. 2018; Dhillon et al. 2021). By utilizing open-source software frameworks such as Python, the system can be modified and extended to incorporate additional functionalities or adapt to new experimental protocols.

Our Python-based software stack includes libraries for real-time data acquisition, signal processing, and feedback control. For instance, we utilize OpenCV for video processing and DeepLabCut-Live (A. Mathis et al. 2018; Kane et al. 2020) for pose estimation, which are essential components in experiments requiring precise monitoring and feedback based on animal behavior. Additionally, we have integrated libraries for handling neural data streams, such as NumPy and SciPy, which facilitate the implementation of complex experimental designs (Akam et al. 2022) involving multiple data sources and feedback modalities. By integrating the system with LED drivers and opsin-expressing transgenic mouse lines, it is straightforward to achieve precise temporal control over neural activation, enabling the study of causal relationships between neural circuit activity and behavior (Lee et al. 2020).

The cost-effectiveness of our system is a significant advantage, making it accessible to a broader range of research labs, including those with limited funding. The use of off-the-shelf components and open-source software drastically reduces the overall cost compared to commercially available systems, which often require expensive proprietary hardware and software licenses (C. Allen and Mehler 2019; White et al. 2019). The open-source nature of our closed-loop platform fosters collaboration and innovation within the research community. By providing unrestricted access, other labs can freely modify, enhance, and share the system, promoting a culture of transparency and shared progress. Importantly, our goal is to increase the number of studies utilizing this platform, enabling a broader exploration of closed-loop methodologies across diverse research contexts. This approach not only accelerates the pace of discovery but also enhances reproducibility and adaptability, ensuring the platform remains a robust tool for advancing neuroscience research.

The system’s flexibility is demonstrated by its successful application across two closed-loop experimental paradigms (CLNF, CLMF). For example, in our study, we utilized the system to implement real-time feedback based on intracellular calcium-induced fluorescence imaging in awake, behaving mice. The system provided auditory feedback in response to changes in cortical activation, allowing us to explore the role of real-time feedback in modulating both neural activity and behavior.

### Importance of closed-loop feedback systems

Closed-loop feedback systems have gained recognition for their ability to modulate neural circuits and behavior in real time, an approach that aligns with the principles of motor learning and neuroplasticity. The ability of mice to learn and adapt to tasks based on cortical or behavioral feedback underscores the parallels between closed-loop systems and natural proprioceptive mechanisms, where continuous sensory feedback is crucial for motor coordination and spatial awareness. This study builds upon the foundational work of (Fetz 1969), who first explored the potential of closed-loop neurofeedback, and expands its application to modern neuroscience by incorporating optical brain-computer interfaces.

Our findings are consistent with previous research showing that rodents can achieve volitional control over externally represented variables linked to their behavior (Clancy et al. 2014; Neely, Piech, et al. 2018; Prsa, Galiñanes, and Huber 2017; Clancy and Mrsic-Flogel 2021). Moreover, the rapid adaptation observed in mice when task rules were altered demonstrates the system’s capacity to facilitate learning and neuroplasticity, even when the conditions for achieving rewards are modified. The quick recovery of task performance after a rule change, as evidenced by the improved performance within days of training, highlights the robustness of the closed-loop feedback mechanism. We have focused on large regional cortical GCAMP signals that are relatively slow in kinetics.

### Neuroplasticity and cortical dynamics

The observation that cortical responses became more localized as mice learned the rewarded behavior aligns with established theories of motor learning, where neural efficiency improves with practice (Makino et al. 2016; Huber et al. 2012; C. B. Allen, Celikel, and Feldman 2003). As mice became more proficient in the task, the widespread cortical activity observed during the initial training sessions became more regionally localized, indicating more efficient neural processing. The reduction in ΔF/F_0_peak values across sessions suggests that the brain becomes more efficient at processing task-relevant information, a phenomenon consistent with the optimization of neural circuits observed in skilled motor learning (Wolpert, Diedrichsen, and Flanagan 2011; Krakauer et al. 2019).

The distinct spatiotemporal patterns of cortical activation observed in mice receiving closed-loop feedback further support the role of real-time feedback in enhancing cortical plasticity. The pronounced segregation of task-related and reward-related cortical dynamics in the later training sessions indicates that closed-loop feedback facilitates the refinement of neural circuits involved in motor learning. These findings are in line with previous studies that have demonstrated the importance of temporally precise feedback in accelerating cortical reorganization and enhancing learning outcomes (Ganguly and Carmena 2009).

### Value of the CLoPy platform in systems neuroscience

The CLoPy platform addresses a critical gap in systems neuroscience by providing a flexible and cost-effective closed-loop feedback system tailored to modulate neural activity and behavior in real time. Many current neuroscience studies assess brain activity and behavior separately, with analyses performed post hoc. CLoPy advances this paradigm by enabling researchers to directly investigate the dynamic interplay between these domains through real-time feedback, a fundamental principle of neuroplasticity and motor learning (Fetz 1969; Wolpert, Diedrichsen, and Flanagan 2011). This integration has broad implications for experimental neuroscience, allowing researchers to probe causal relationships between brain activity, behavior, and environmental contingencies in a controlled and scalable manner.

By enabling closed-loop experiments based on neural imaging (e.g., GCaMP fluorescence) and real-time behavioral tracking (e.g., DeepLabCut-Live), CLoPy allows researchers to address a range of experimental questions. For example, the system can be used to study how cortical plasticity facilitates motor learning, how the brain adapts to rule changes in a task, and how feedback mechanisms influence recovery from neurological injuries. These capabilities align with the needs of systems neuroscience researchers who seek to elucidate the mechanisms underlying adaptive brain-behavior interactions across different experimental conditions. Compared to other platforms, such as pyControl (Akam et al. 2022), CLoPy uniquely integrates mesoscopic calcium imaging, real-time tracking, and feedback delivery, making it particularly suitable for exploring neural dynamics at a finer resolution.

### Positioning within the open-source neuroscience community

As part of the broader open-source neuroscience movement, CLoPy exemplifies the principles of accessibility, modularity, and reproducibility. The platform’s reliance on widely available hardware, such as Raspberry Pi and Nvidia Jetson, and its use of Python-based libraries ensure that researchers with varying levels of technical expertise can adopt and modify it. This is particularly important in the current landscape, where open-source tools like pyControl (Akam et al. 2022), OpenEphys (Siegle et al. 2017), and Bonsai (Lopes and Monteiro 2021) have transformed experimental neuroscience by reducing costs and fostering collaboration.

CLoPy complements these efforts by extending open-source frameworks to the realm of closed-loop neuroscience. It provides detailed documentation (https://clopy-docs.readthedocs.io), hardware schematics, and adaptable software, empowering researchers to customize the system for diverse experimental needs. For example, CLoPy enables experiments involving optogenetic stimulation, multimodal feedback, and integration with advanced imaging techniques. By lowering barriers to entry, it democratizes access to closed-loop experimental paradigms, particularly for labs with limited resources.

### Experimental applications and implications

CLoPy’s flexibility and modularity make it suitable for a wide range of experimental applications. In this study, we demonstrated its utility in closed-loop neural feedback (CLNF) and closed-loop movement feedback (CLMF) paradigms. These experiments showcase how CLoPy can be used to study fundamental questions about the dynamic interplay between neural activity, feedback, and behavior. For instance, the system’s ability to deliver graded auditory feedback in response to cortical activation allowed us to explore how real-time neurofeedback modulates neural dynamics. Similarly, its real-time tracking capabilities enabled precise feedback delivery based on specific behavioral events, such as limb movements, facilitating the investigation of motor learning and adaptation. In our system, we measured latencies of ∼63 ms for CLNF and ∼67 ms for CLMF. While such latencies may limit applications requiring millisecond precision, such as fast whisker movements, saccades, or fine-reaching kinematics, we emphasize that many relevant behaviors, including postural adjustments, limb movements, locomotion, and sustained cortical state changes, occur on timescales that are well within the capture range of our system. It is also important to note that these latencies are not solely dictated by hardware constraints. A significant component arises from the inherent biological dynamics of the calcium indicator (GCaMP6s) and calcium signaling itself, which introduce slower temporal kinetics independent of processing delays. Newer variants, such as GCaMP8f, offer faster response times and could further reduce effective biological latency in future implementations. We acknowledge that Raspberry Pi provides a low-cost solution but contributes to modest computational delays, while Nvidia Jetson offers faster inference at higher cost. Our choice reflects a balance between accessibility, cost-effectiveness, and performance, making the system deployable in many laboratories. Importantly, the modular and open-source design means the pipeline can readily be adapted to higher-performance GPUs or integrated with electrophysiological recordings, which provide higher temporal resolution. In the case of CLNF we have focused on large regional cortical GCAMP signals that are relatively slow in kinetics. While such changes are well suited for transcranial mesoscale imaging assessment, it is possible that cellular 2-photon imaging (Yu et al. 2021) or preparations that employ cleared crystal skulls (Kim et al. 2016) could resolve more localized and higher frequency kinetic signatures. In this case it would be important to potentially tune and enhance closed-loop methodology to more closely follow these events.

The platform’s ability to modulate neural and behavioral outputs in real time has far-reaching implications for experimental neuroscience. It enables researchers to study neural mechanisms underlying learning, adaptation, and plasticity with a level of precision and scalability that was previously difficult to achieve. For example, CLoPy can be used to investigate how the brain reorganizes itself after injury or how neural circuits adapt to changing task demands.

### Future directions and broad impact

To further enhance its utility, future iterations of CLoPy could integrate advanced machine learning algorithms for real-time data analysis and decision-making, enabling more sophisticated feedback paradigms. Additionally, expanding the platform’s compatibility with other experimental modalities, such as multi-photon imaging and electrophysiology, could broaden its applications in neuroscience research. CLoPy’s scalability also opens avenues for its use in translational research, such as testing neurorehabilitation interventions or brain-computer interfaces in larger animal models or humans. By providing an open-source, flexible, and cost-effective solution, CLoPy has the potential to significantly advance the field of systems neuroscience. It not only enables researchers to address complex experimental questions but also fosters collaboration and innovation within the neuroscience community. As more labs adopt and refine this platform, we anticipate that it will accelerate the pace of discovery, enhance reproducibility, and contribute to the development of novel therapeutic strategies for neurological disorders. While we aimed to include multiple animals for each experimental condition, certain task rules are represented by only one or a few animals. This limitation primarily arises from logistical challenges, including the extensive time required for training under specific task contingencies and the variability in maintaining high-quality optical windows over prolonged experimental periods. Consequently, while these data provide valuable insights into the feasibility and mechanisms of the task, we acknowledge that broader generalizations should be made cautiously and that further replication will be important in future studies.

In conclusion, our study highlights the significant potential of closed-loop feedback systems for advancing neuroscience research. By providing a flexible, cost-effective, and open-source platform, we offer a valuable tool for exploring the complex interactions between brain activity and behavior, with implications for both basic research and clinical applications.

## Materials and methods

### Animals

Mouse protocols were approved by the University of British Columbia Animal Care Committee (ACC) and followed the Canadian Council on Animal Care and Use guidelines (protocol A22-0054). A total of 56 mice (postnatal 104-140) were used in this study: 26 female and 30 male transgenic C57BL/6 mice expressing GCaMP6s were used. CLNF experiments (n=40, 17 females, 23 males) were done with tetO-GCaMP6s x CAMK tTA (Wekselblatt et al., 2016), and CLMF experiments (n=16, 9 females, 7 males) were done with Ai94 from the Allen Institute for Brain Science, crossed to Emx1–cre and CaMK2-tTA line (Jackson Labs) (Madisen et al., 2015). Mice were housed in a conventional facility in plastic cages with micro-isolator tops and kept on a normal 12 hr. light cycle with lights on at 7 AM. Most experiments were performed toward the end of the mouse light cycle. Mice that were unable to achieve a success rate of 70% after 7 days of training in CLNF experiments were excluded from the study (total 6 mice).

### Animal surgery, chronic transcranial window preparation

Animals were anesthetized with isoflurane, and a transcranial window was installed as previously described (Silasi et al. 2016; Vanni and Murphy 2014) and in an amended and more extensive protocol described here. A sterile field was created by placing a surgical drape over the previously cleaned surgical table, and surgical instruments were sterilized with a hot bead sterilizer for 20 s (Fine Science Tools; Model 18000–45). Mice were anesthetized with isoflurane (2% induction, 1.5% maintenance in air) and then mounted in a stereotactic frame with the skull level between lambda and bregma. The eyes were treated with eye lubricant (Lacrilube; www.well.ca) to keep the cornea moist, and body temperature was maintained at 37°C using a feedback-regulated heating pad monitored by a rectal probe. Lidocaine (0.1 ml, 0.2%) was injected under the scalp, and mice also received a 0.5 ml subcutaneous injection of a saline solution containing buprenorphine (2 mg/ml), atropine (3 μg/ml), and glucose (20 mM). The fur on the head of the mouse (from the cerebellar plate to near the eyes) was removed using a fine battery-powered beard trimmer, and the skin was prepared with a triple scrub of 0.1% Betadine in water followed by 70% ethanol. Respiration rate and response to toe pinch were checked every 10–15 min to maintain the surgical anesthetic plane.

Before starting the surgery, a cover glass was cut with a diamond pen (Thorlabs, Newton, NJ, USA; Cat#: S90W) to the size of the final cranial window (∼9 mm diameter). A skin flap extending over both hemispheres approximately 3 mm anterior to bregma and to the posterior end of the skull and down lateral was cut and removed. A #10 scalpel (curved) and sterile cotton tips were used to gently wipe off any fascia or connective tissue on the skull surface, making sure it was completely clear of debris and dry before proceeding. The clear version of C and B-Metabond (Parkell, Edgewood, NY, USA; Product: C and B Metabond) dental cement was prepared by mixing 1 scoop of C and B Metabond powder (Product: S399), 7 drops of C and B Metabond Quick Base (Product: S398), and one drop of C and B Universal catalyst (Product: S371) in a ceramic or glass dish (do not use plastic). Once the mixture reaches a consistency that makes it stick to the end of a wooden stir stick, a titanium fixation bar (22.2 × 2.7 × 3.2 mm) was placed so that there was a 4 mm posterior space between the bar edge and bregma, by applying a small amount of dental cement and holding it pressed against the skull until the cement partially dried (1–2 min). With the bar in place, a layer of dental adhesive was applied directly on the intact skull. The precut cover glass was gently placed on top of the mixture before it solidified (within 1 min), taking care to avoid bubble formation. If necessary, extra dental cement was applied around the edge of the cover slip to ensure that all the exposed bone was covered, and that the incision site was sealed at the edges. The skin naturally tightens itself around the craniotomy, and sutures are not necessary. The mixture remains transparent after it solidifies, and one should be able to clearly see large surface veins and arteries at the end of the procedure. Once the dental cement around the coverslip is completely solidified (up to 20 min), the animal received a second subcutaneous injection of saline (0.5 ml) with 20 mM of glucose and was allowed to recover in the home cage with an overhead heat lamp and intermittent monitoring (hourly for the first 4 hr. and every 4–8 hr. thereafter for activity level). Then, the mouse was allowed to recover for 7 days before task training.

### Water deprivation and habituation to experiments

Around 10–21 days after the surgery, the animals were placed on a schedule of water deprivation. Given the variation in weight due to initial ad libitum water consumption, the mouse weight was defined 24 hr. after the start of water restriction. If mice did not progress well through training, they were still given up to 1 ml of water daily, there was also a 15% maximal weight loss criterion used for supplementation (see detailed protocol). All mice were habituated for 5 days before data collection. Awake mice were head-fixed and placed in a dark imaging chamber for training and data collection for each session. We tried to not keep animals headfixed for more than 45 minutes in each session as they become less engaged with long duration headfixed sessions. After headfixing them, it takes about 15 minutes to get the experiment going and therefore 30 - 40 minutes long recorded sessions seemed appropriate before they stop being engaged or before they get satiated in the task. High performing animals were able to maintain body weight and gain weight towards pre-surgery and pre-water restriction values.

### CLNF, CLMF setup and behavior experiments

Our goal has been to deliver a robust, cross-platform, and cost-effective solution for closed-loop feedback experiments. We have designed and tested a behavioral paradigm where head-fixed mice learn an association between their cortical or behavioral activity, external feedback, and rewards. We tested our CLNF system on Raspberry Pi, and CLMF system on an Nvidia Jetson GPU device, for their compactness, general-purpose input/output (GPIO) programmability, and wide community support. While our investigations center around mesoscale cortical imaging of genetically encoded calcium sensors and behavior imaging, the approach could be adapted to any video or microscopy-dependent signal where relative changes in brightness or keypoint behavior are observed and feedback is given based on a specific rule. This system benefits from advances in pose estimation (Forys et al. 2020; Kane et al. 2020)and employs strategies to improve real-time processing of pre-defined keypoints on compact computers such as the Nvidia Jetson. We have constructed a software and hardware-based platform built around open-source components.

To examine the performance of the closed-loop system, we used water-restricted adult transgenic mice that expressed GCaMP6s widely in the cortex (details in the methods section and in Table 5 and Table 6). Using the transcranial window imaging technique, we assessed the ability of these animals to control brain regions of interest and obtain water rewards. After an initial period of habituation to manual head fixation, mice were switched to closed-loop task training. In training, we had two primary groups: 1) a single cortical ROI linked to water reward and the subject of auditory feedback (Animation 1A, 1B, 1C), or 2) a pairing between two cortical ROIs where auditory feedback and water rewards were given based on the difference in activity between sites (Animation 1D, 1E, 1F).

For neurofeedback-CLNF, we calculate an online ΔF/F_0_ value (GCAMP signal) which represents a relative change of intensity in the region(s) of interest. In brief, a running baseline (F_0_) is computed at each sample point, subtracted and divided (ΔF/F_0_) from the raw fluorescence values; more details about online ΔF/F_0_ calculation are in the methods section. These calculations are made in near real-time on the Raspberry Pi and are used to control the GPIO output pins that provide digital signals for closed-loop feedback and water rewards to water-restricted mice. In 1-ROI experiments, we mapped the range of average ΔF/F_0_ values in the ROI (Figure 1) to a range of audio frequencies (1 kHz - 22 kHz), which acted as feedback to the animal. For 2-ROI experiments, the magnitude of the ΔF/F_0_ activity difference between the ROIs, based on the specified rule (e.g., ROI1-ROI2), was mapped to the range of audio frequencies. We confirmed that these frequencies were accurately generated and mapped by audio recordings (Supplementary Figure 2 and see Methods) obtained at 200 kHz using an ultrasonic microphone (Dodotronic, Ultramic UM200K) positioned within the recording chamber ∼10 cm from the audio speaker. To confirm the timing of feedback latency, LED lights were triggered instead of water rewards (Supplementary Figure 1), and the delay was calculated between the detected event (green LED ON for CLNF and paw movement for CLMF) and the red LED flash. In the case of CLNF, the camera recording brain activity was used to record both the flashing green and red LEDs. Temporal traces of green and red LED pixels were extracted from the recorded video, and the average delay between the green and red LEDs becoming bright was calculated as the delay in closed-loop feedback for CLNF experiments. Performing this analysis indicated that the Raspberry Pi system could provide reliable graded feedback using GPIO within ∼63 ± 15 ms for CLNF experiments.

An imaging rig was developed using two Raspberry Pi 4B+ single-board computers, designated as “brain-pi” (master device for initiating all recordings) for widefield GCaMP imaging and “behavior-pi” (slave device waiting for a trigger to start recording) for simultaneous behavior recording. Both devices were connected to the internet via Ethernet cables and communicated with each other through 3.3 V transistor-transistor logic (TTL) via general-purpose input outputs (GPIOs). To synchronize session initiation, GPIO pin #17 on the brain-pi (configured as an output) was connected to GPIO pin #17 on the behavior-pi (configured as an input), allowing the brain-pi to send a TTL signal to the behavior-pi at the start of each session. Additionally, frames were aligned based on a LED ON event for each session.

Additional hardware components were integrated into the brain-pi setup. GPIO pin#27 (output) was connected to a solenoid (Gems Sensor, 45M6131) circuit to deliver water rewards, while GPIO pin#12 (output) was linked to a buzzer (Adafruit product #1739) positioned under the head-fixing apparatus to signal trial failures. GPIO pin#21 (output) was used to trigger a LED driver controlling both short blue (447.5 nm) and long blue (470 nm) LEDs, which were essential for cortical imaging. A speaker was connected to the brain-pi’s 3.5 mm audio jack to provide auditory output (PulseAudio driver) during the experiments. Both the brain-pi and behavior-pi devices were equipped with compatible external hard drives, connected via USB 3.0 ports, to store imaging and behavioral data, ensuring reliable data capture throughout the experimental sessions.

Mice with implanted transcranial windows on the dorsal cortex were head-fixed in a transparent acrylic tube (1.5-inch outer diameter, 1 ⅛ inch inner diameter) and placed such that the imaging camera (RGB Raspberry Pi Camera, OmniVision OV5647 CMOS sensor) was above the transcranial window, optimally focused on the cortical surface for GCaMP imaging. The GCaMP imaging cameras had lenses with a focal length of 3.6 mm and a field of view of ∼10.2 × 10.2 mm, leading to a pixel size of ∼40 μm, and were equipped with triple-bandpass filters (<u>Chroma 69013m</u>), which allowed for the separation of GCaMP epifluorescence signals and 447 nm reflectance signals into the green and blue channels, respectively. The depth of field (∼3 mm) was similar to previous reports (Lim et al. 2012) which provided both a large focal volume over which to collect fluorescence and made the system less sensitive to potential changes in the z-axis position. To reduce image file size, we binned data at 256 × 256 pixels on the camera for brain imaging data. Brain images were saved as 8-bit RGB stacks in HDF5 file format. We manually fixed the camera frame rate to 15 Hz, turned off automatic exposure and auto white balance, and set white balance gains to unity. For both green epifluorescence and blue reflection channels, we adjusted the intensity of illumination so that all values were below 180 out of 256 grey levels (higher levels increase the chance of cross-talk and saturation).

Mesoscale GCaMP imaging can often be performed with single-wavelength illumination (Gilad and Helmchen 2020; Nakai et al. 2023). However, in this experiment, we utilized dual-LED illumination of the cortex (Michelson et al. 2023). One LED (short-wavelength blue, 447 nm Royal Blue Luxeon Rebel LED SP-01-V4 paired with a Thorlabs FB 440-10 nm bandpass filter) monitored light reflectance to account for hemodynamic changes (Xiao et al. 2017), while the second LED (long-wavelength blue, 470 nm Luxeon Rebel LED SP-01-B6 combined with a Chroma 480/30 nm filter) was used to excite GCaMP for green epifluorescence. Both signals were captured simultaneously using an RGB camera. For each mouse, light from both the excitation and reflectance LEDs was channeled into a single liquid light guide, positioned to illuminate the cortex (Figure 1v) (Michelson et al. 2023). Further specifics are outlined in the accompanying Parts List and assembly instructions document. A custom-built LED driver, controlled by a Raspberry Pi, activated each LED at the beginning of the session and deactivated them at the session’s end. This on-off illumination shift was later used in post hoc analysis to synchronize frames from the brain and behavior cameras.

Before initiating the experimental session, key session parameters were configured in the config.ini file, as detailed in the “Key Configuration Parameters” section below. Additionally, we ensured that the waterspout was positioned appropriately for the mouse to access and consume the reward. To launch the experiment, open a terminal on the brain-pi (master) device, ensuring that the current directory is set to clopy/brain/. The experiment can be started by entering the command python3 <script_name>.py, where <script_name> corresponds to the appropriate Python script for the session (Supplementary Figure 7). We provide two pre-defined scripts in the codebase: one for 1ROI and another for 2ROI experiments. Both scripts are functionally similar. Upon execution, the script prompts for the “mouse_id,” which can be entered as an array of characters, followed by pressing ‘Enter.’ Upon initialization, two preview windows appear. The first window displays live captured images with intensity values overlaid in the green and blue channels, which allow for adjustments to LED brightness levels. The second window displays real-time ΔF/F_0_ values overlaid on cortical locations, enabling brain alignment checks. These preview windows are used to assess imaging quality and ensure appropriate settings before starting the experiment (Supplementary Figure 6). Once all settings are confirmed, pressing the ‘Esc’ key starts the experiment. At this point, only one window displaying incoming images is shown. The experiment begins with a rest period of duration specified in the config.ini file, followed by alternating trial and rest periods. Data acquired during the session are saved in real-time on the device. For more details and the latest instructions on system setup, please refer to the project’s GitHub page - https://github.com/pankajkgupta/clopy, and documentation - https://clopy-docs.readthedocs.io.

An additional imaging rig for CLMF was developed utilizing an Nvidia-Jetson Orin device (8-core ARM Cortex CPU, 2048 CUDA cores, 64 GB memory), which served as the master device responsible for triggering all recordings. The Nvidia-Jetson was connected to an Omron Sentech STC-MCCM401U3V USB3 Vision camera for behavior imaging and real-time pose tracking. A personal computer (serving as the slave device) running EPIX software and frame grabber (PIXCI® E4 Camera Link Frame Grabber) was connected to a Pantera TF 1M60 CCD camera (Dalsa) for widefield GCaMP imaging. Both the Nvidia-Jetson and the EPIX PC were linked via transistor-transistor logic (TTL) for communication. For session synchronization, the GPIO pin #17 on the Nvidia-Jetson, configured as an output, was connected to the trigger input pin of the EPIX framegrabber E4DB. This configuration enabled the Nvidia-Jetson to send a TTL signal to initiate frame capture on the EPIX system at the start of each session. Additionally, this same pin was connected to an LED driver to trigger the activation of a long blue LED (470 nm) for GCaMP imaging simultaneously.

Several hardware components were integrated into the Nvidia-Jetson setup to support experimental protocols. GPIO pin #13 (output) was linked to a solenoid circuit to deliver water rewards, while GPIO pin #7 (output) was connected to a buzzer placed under the head-fixation apparatus to signal trial failures. Auditory feedback during the experiments was provided through a speaker connected to the Nvidia-Jetson’s audio output. Both the Nvidia-Jetson and EPIX PC were equipped with sufficient storage space to ensure the reliable capture and storage of behavioral video recordings and widefield image stacks, respectively, throughout the experimental sessions.

Mice, implanted with a transcranial window over the dorsal cortex, were head-fixed in a custom-made transparent, rectangular chamber (dimensions provided). The chamber was designed using CAD software and fabricated from a 3 mm-thick acrylic sheet via laser cutting. Complete model files, acrylic sheet specifications, and assembly instructions are available on the GitHub repository. A mirror was positioned at the bottom and front of the chamber (Figure 1B) to allow multiple views of the mouse for improved tracking accuracy. The Omron Sentech camera was equipped with an infrared (IR)-only filter to exclusively capture IR illumination, effectively blocking other light sources and ensuring consistent behavioral imaging for accurate pose tracking. For widefield GCaMP imaging, the Dalsa camera was equipped with two front-to-front lenses (50 mm, f/1.435 mm and f/2; Nikon Nikkor) and a bandpass emission filter (525/36 nm, Chroma). The 12-bit images were captured at a frame rate of 30 Hz (exposure time of 33.3 ms) with 8×8 on-chip spatial binning (resulting in 128×128 pixels) using EPIX XCAP v3.8 imaging software. The cortex was illuminated using a blue LED (473 nm, Thorlabs) with a bandpass filter (467–499 nm) to excite calcium indicators, and the blue LED was synchronized with frame acquisition via TTL signaling. GCaMP fluorescence image stacks were automatically saved to disk upon session completion.

Prior to starting a session, key experimental parameters were configured within the config.ini file (see “Key Configuration Parameters” below for details). The waterspout was positioned close to the mouse to allow easy access for licking and consuming rewards. To initiate the experiment on the Nvidia-Jetson (master device), the terminal’s current directory was set to clopy/behavior/. The experiment was launched by executing the command python3 <script_name>.py, where <script_name> corresponds to the Python script responsible for running the experiment (Supplementary Figure 6). The provided script maps the speed of a specific control-point (FLL_bottom) to the audio feedback and executes 60 trials. Once launched, the program prompts for the “mouse_id,” which could be entered as an array of characters, followed by pressing Enter. A preview window displaying a live behavioral view was then presented, allowing for adjustments of IR brightness levels and the waterspout (Supplementary Figure 6). Once imaging quality and settings were confirmed to be optimal, pressing the “esc” key started the experiment. During the experiment, the tracked points were overlaid on the real-time video, and the session alternated between rest and trial periods. All acquired data were saved locally on the Nvidia-Jetson during the experiment and automatically transferred to the EPIX PC after completion. Additional setup instructions and system details can be found on the associated GitHub page and documentation - https://clopy-docs.readthedocs.io.

### Key configuration parameters

Prior to running the experiment script, session parameters were set in the config.ini file under a designated configuration section. This file contains multiple config sections, each tailored to a different experiment type, such as brain-pi or behavior-pi. The appropriate config section is specified in the experiment script. Key parameters in this section are listed in the table below.

Further details on these parameters can be accessed via the GitHub repository and the documentation at https://clopy-docs.readthedocs.io.

### CLoPy platform

Closed-Loop Feedback Training System (CLoPy) is an open-source software and hardware system implemented in the Python (>=3.8) programming language. This work is accompanied by a package to replicate the system, reproduce figures in this publication, and an extensive supplemental guide with full construction illustrations and parts lists to build the platform used. See https://github.com/pankajkgupta/clopy and https://clopy-docs.readthedocs.io for details and accompanying acquisition and analysis code. We also provide model files for machined and 3D-printed parts in the repository; links to neural and behavioral data can also be found at the federated research data repository- https://doi.org/10.20383/102.0400.

While the presented CLNF experiments were conducted on a Raspberry Pi 4B+ device, the system can be used on any other platform where the Python runtime environment is supported. Similarly, CLMF experiments were conducted on an Nvidia Jetson Orin device, but it can be deployed on any other device with a GPU for real-time inference. For any programmable camera to be used with the system, one can implement a wrapper Python class that implements the CameraFactory interface functions for integration with the system.

### Graded feedback (online ΔF/F_0_-audio mapping)

The auditory feedback was proportional to the magnitude of the neural activity. We translated fluorescence levels into the appropriate feedback frequency and played the frequency on speakers mounted on two sides of the imaging platform. Frequencies used for auditory feedback ranged from 1 to 22 kHz in quarter-octave increments (Han et al. 2007; Clancy et al. 2014). When a target was hit, a circuit driven solenoid delivered a reward to mice.

To map the fluorescence activity *F* to the quarter-octave index *n*, we used linear scaling. Let *F*_*min*_ and *F*_*max*_ be the minimum and maximum fluorescence values.*n*_*max*_ The maximum number of quarter-octave steps between 1 kHz and 22 kHz.

The number of quarter-octave steps between 1 kHz and 22 kHz can be calculated as:

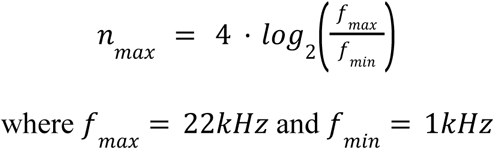

The quarter-octave index *n* can be mapped linearly from fluorescence activity *F* as:

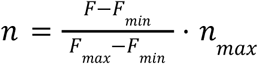

Thus, the final equation mapping fluorescence activity to audio frequency is:

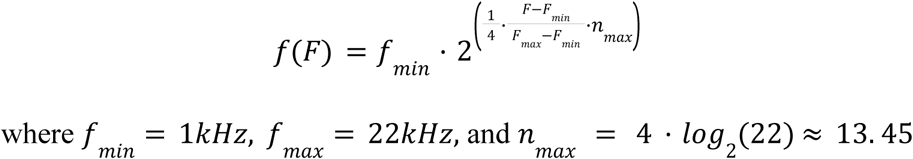

This equation allows us to map any fluorescence activity *F* within the range [*F*_*min*_, *F*_*max*_] to a corresponding frequency within the 1–22 kHz range, in quarter-octave increments.

### Checking the dynamic range of graded auditory feedback

To assess the dynamic range of audio signals from the speakers, we used an Ultra microphone (Dodotronic) to record feedback signals generated during a session, detect the frequencies, and compare the detected frequencies through our speakers to what was commanded through the program. We verified a linear relationship between 1 and 22 kilohertz (Supplementary figure 2). Analysis of GCAMP imaging experiments indicated that baseline activity values were associated with mapped sounds in the 6±4 kHz range. At reward points, which are threshold-dependent (Delta F over F values), the commanded auditory feedback values were significantly higher (18±2 kHz range). A previous version of the CLNF system was found to have non-linear audio generation above 10 kHz, partly due to problems in the audio generation library and partly due to the consumer-grade speaker hardware we were employing. This was fixed by switching to the Audiostream (https://github.com/kivy/audiostream) library for audio generation and testing the speakers to make sure they could output the commanded frequencies (supplementary figure 2).

### Determining reward threshold based on a baseline session

Before starting the experiments, and after the habituation period, a baseline session (day 0) of the same duration as the experiments is recorded. This session is similar to the experimental sessions in the coming days, except that the mice do not receive any rewards. Offline analysis of this session is used to establish a threshold value for the target activity (see the target activity section to read more). In brief, for 1 ROI experiments, target activity is the average ΔF/F_0_ activity in that ROI. For 2 ROI experiments, target activity is based on the specified rule in the config.ini file. For example, target activity for the rule “ROI1-ROI2” would be “average ΔF/F_0_ activity in ROI1 - average ΔF/F_0_ activity in ROI2.”

### Determining ROI(s) for changes in CLNF task rules

Dorsal cortical widefield activity is dynamic, and ongoing spatiotemporal motifs involve multiple regions changing activity. Choosing ROI(s) for CLNF has some caveats and requires some considerations before choosing. It was relatively straightforward for initial training experiments where baseline (day 0) sessions were used to establish a threshold for the selected ROI on day 1. Mice learn to modulate the selected ROI over the training sessions. Interestingly, other cortical ROIs were also changing along with the target ROI as mice were learning the task. We needed to be careful when changing the ROI on day 11 because if we changed to an ROI that also changes along with the previous ROI, mice could keep getting rewards without realizing any change. To address this issue, we analyzed the neural data from day 1 to day 10 and found potential ROIs for which the threshold crossings did not increase significantly or were not on par with the previous ROI, and one of these ROIs was chosen for the rule change.

### Online ΔF/F_0_ calculation

In calcium imaging, ΔF/F_0_ is often used to represent the change in fluorescence relative to a baseline fluorescence (F_0_), which helps in normalizing the data. For CLNF real-time feedback, we computed ΔF/F_0_ online, frame by frame.

Let *F*(*t*) represent the fluorescence signal at time.*t*. A running baseline *F*_0_(*t*)is estimated by applying a sliding window to the fluorescence signal to calculate a moving average over a window of size *N*.

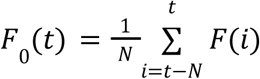

Once the running baseline.*F*_0_(*t*)is computed, the ΔF/F₀ at time *t* is calculated as:

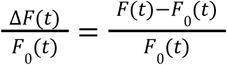

Each incoming frame contained both a green channel (capturing GCaMP6s fluorescence) and a short blue channel (representing blood volume reflectance) (Ma et al. 2016; Wekselblatt et al. 2016; Valley et al. 2020). These frames were appended to a doubly-ended queue (deque), a Python data structure, for running baseline *F*_0_(*t*)The length of the deque.*N* was defined by the product of two parameters from the configuration file: “dff_history” (in seconds) and “framerate” (in frames per second), which determined the number of frames used for the running baseline. For the CLNF experiments, we used a dff_history of 6 s (a dynamic baseline) and a framerate of 15 fps, resulting in a deque length of 90 frames. When the deque reached full capacity, appending a new frame automatically removed the oldest frame, ensuring an updated running ΔF/F_0_ throughout the experiment, regardless of session duration. This method effectively avoided memory limitations over time. We would like to emphasize that we are not directly averaging activity over 6 s to compare against the reward threshold. Instead, the preceding 6 s of activity is used solely to compute a dynamic baseline for ΔF/F_0_ (ΔF/F_0_ = (F – F_0_)/F_0_). Here, F_0_ is calculated as the mean fluorescence intensity over the prior 6 s window and is updated continuously throughout the session. This baseline is then subtracted from the instantaneous fluorescence signal to detect relative changes in activity. The reward threshold is therefore evaluated against these baseline-corrected ΔF/F_0_ values at the current time point, not against an average over 6 s. This moving-window baseline correction is a standard approach in calcium imaging analyses, as it helps control for slow drifts in signal intensity, bleaching effects, or ongoing fluctuations unrelated to the behavior of interest. Thus, the 6 s window is not introducing a temporal lag in reward assignment but is instead providing a reference to detect rapid increases in cortical activity. For each channel (green and blue), the running F_0_ was subtracted from each incoming frame to obtain ΔF, followed by ΔF/F_0_ calculations, applied per pixel as a vectorized operation in Python. To correct for hemodynamic artifacts (Ma et al. 2016), we subtracted the blue excitation and epifluorescence channel ΔF/F_0_ from the green channel ΔF/F_0_.

It is important to note that while blood volume reflectance is typically captured using green light (Ma et al. 2016), we used short blue light due to technical constraints associated with the Raspberry Pi camera’s rolling shutter which made strobing infeasible. The short blue light (447 nm) with a 440 ± 5 nm filter is close to the hemoglobin isosbestic point and has been shown to correlate well with the 530 nm green light signal as a proxy for hemodynamic activity (Michelson et al. 2023; Murphy et al. 2020). Additionally, the 447 nm LED would be expected to produce minimal green epifluorescence at the low power settings used in our experiments (Dana et al. 2014). Previous studies have evaluated and compared the performance of corrected versus uncorrected signals using this method (Xiao et al. 2017; Murphy et al. 2020).

### Offline ΔF/F_0_ calculation

For offline analysis of GCaMP6s fluorescence in CLNF experiments, the green and blue channels (Ma et al. 2016; Wekselblatt et al. 2016; Valley et al. 2020) were converted to ΔF/F_0_ values. For each channel, a baseline image (F_0_) was computed by averaging across all frames of the recording session. The F_0_ was then subtracted from each individual frame producing a difference image (ΔF). This difference was divided by F_0_, resulting in the fractional change in intensity (ΔF/F_0_) for each pixel as a function of time. To further correct for hemodynamic artifacts (Ma et al., 2016), the blue channel reflected light ΔF/F_0_ signal, reflecting blood volume changes, was subtracted from the green channel ΔF/F_0_ signal, isolating the corrected GCaMP6s fluorescence response from any potential confounding vascular contributions.

Offline analysis of GCaMP6s fluorescence in CLMF experiments involved processing the green epifluorescence channel. We did not collect the hemodynamic signal in this experiment because we intended to employ the second channel for optogenetic stimulation (not included in this article). A baseline image (F_0_) was computed by averaging across all frames of the recording session. The F_0_ was then subtracted from each individual frame, producing a difference image (ΔF). This difference was divided by F_0_, resulting in the fractional change in intensity (ΔF/F_0_) for each pixel as a function of time.

### Seed-pixel correlation matrices

Widefield cortical image stacks were registered to the Allen Mouse Brain Atlas (Wang et al. 2020; Chon et al. 2019) and segmented into distinct regions, including the olfactory bulb (OB), anterolateral motor cortex (ALM), primary motor cortex (M1), secondary motor cortex (M2), sensory forelimb (FL), sensory hind limb (HL), barrel cortex (BC), primary visual cortex (V1), and retrosplenial cortex (RS) in both the left and right cortical hemispheres. The average activity in a 0.4 x 0.4 mm² area (equivalent to a 10 x 10 pixel region) centered on these regions (also referred to as seed pixels) was calculated, representing the activity within each region. These signals were then used to generate temporal plots. The temporal plots were epoched into trial and rest conditions, and correlations between the regions were computed, resulting in correlation matrices for each condition across each day. Each element of the matrix represents the pairwise correlation between two cortical regions. For each pair of cortical regions, we obtained correlation values during both trial and rest periods for every session, creating a time series (over sessions) with two conditions (trial and rest).

## Supporting information

supplemental animation 1 closed-loop neurofeedback

supplemental animation 2 closed-loop movement feedback forelimb left

supplemental animation 3 closed-loop movement feedback forelimb right

## Acknowledgements

This work was supported by Canadian Institutes of Health Research (CIHR) foundation grant FDN-143209 and project grant PJT-180631 (to T.H.M.). T.H.M. was also supported by the Brain Canada Neurophotonics Platform, a Heart and Stroke Foundation of Canada grant in aid, the Natural Sciences and Engineering Council of Canada (NSERC; GPIN-2022-03723), and a Leducq Foundation grant. This work was supported by resources made available through the Dynamic Brain Circuits cluster and the NeuroImaging and NeuroComputation Centre at the UBC Djavad Mowafaghian Centre for Brain Health (RRID SCR_019086) and made use of the DataBinge forum, and computational resources and services provided by Advanced Research Computing (ARC) at the University of British Columbia. We thank Pumin Wang and Cindy Jiang for surgical assistance, Jamie Boyd and Jeffrey M LeDue for technical assistance.

## Data and code availability

The source data used in this paper is available here- https://doi.org/10.20383/103.01152. Code to replicate the system, recreate the figures, and associated pre-processed data are publicly available and hosted on GitHub - https://github.com/pankajkgupta/clopy. Detailed documentation can be found here-https://clopy-docs.readthedocs.io. Any additional information required to reanalyze the data reported in this work is available from the lead contact upon request.

### Statistics

Various statistical tests were performed to support the analysis presented in accompanying figures. For p-value matrices in Figure 10B, 10C, a two-way repeated-measures ANOVA with Bonferroni post-hoc correction was used to test the significance of correlation changes across two factors: sessions (days 1–10) and condition (trial vs. rest). Significant changes in the correlation matrices along these two variables showed complementary patterns (Figure 10). Changes across sessions involved the bilateral M1, M2, FL, HL, and BC regions, while changes between the trial and rest conditions involved bilateral OB, ALM, V1, and RS regions.

A list of statistical tests used in a figure, its purpose and data used are summarized in the table below.

**Table 2:** List of statistical tests performed.

| Figure | Test name | Purpose | Arguments |
| --- | --- | --- | --- |
| Figure 4A | Mann-Kendall Test<br>(pymannkendall package in Python) | To determine a monotonic upward trend in the success rate of all CLNF mice before the rule change (from day 1 to day 10). | Normalized success rate per mouse before the rule change (N=40) |
|  |  | To determine a monotonic upward trend in the success rate of CLNF rule-change mice after the rule change (day 11 to day 19). | Normalized success rate per mouse after the rule change (N=17) |
| Figure 4A | RM-ANOVA<br>(Pingouin package in Python) | Test the significance of the success rate increase before the rule change in the CLNF No-rule-change group (Day 1 to Day 10). | Normalized success rate per mouse in No-rule-change mice before the rule change (N=23) |
|  |  | Test the significance of the success rate increase after the rule change in the CLNF Rule-change group (day 11 to day 19). | Normalized success rate per mouse in Rule-change mice after the rule change (N=17) |
| Figure 4A | Two way RM-ANOVA<br>(Pingouin package in Python) | Test significance of day and group on the CLNF success rate before rule change (day 1 to day 10) | Normalized success rate per mouse in Rule-change and No-rule-change mice before rule change, along with group labels. |
| Figure 4A | Mann-Whitney U-test with Benjamini-Hochberg correction<br>(pymannkendall package in Python) | Test the difference in the distribution of CLNF success rates between:<br>day 1 no-rule-change and day 3 no-rule-change<br>day 10 No-rule change and day 10 Rule change<br>day 10 rule change and day 11 rule change<br>day 10 rule change and day 19 rule change | Normalized success rate per mouse in Rule-change and No-rule-change mice before rule change, along with group labels. |
| Figure 4B | RM-ANOVA (Pingouin package in Python) | Test the significance of the success rate increase before the rule change in the CLMF Rule-change group (Day 1 to Day 4).<br>Test the significance of the success rate increase after the rule change in the CLMF Rule-change group (from day 5 to day 10).<br>Test the significance of the success rate increase in the CLMF no-rule-change group (day 1 to day 10).<br>Test significance of success rate increase in CLMF No-feedback group (day 1 to day 10). | Normalized success rate per mouse in Rule-change and No-rule-change mice before rule change, along with group labels. |
| Figure 5C | Permutation Test (MMD function in hyppo package) | Multivariate two-sample test to compare ROI1 vs ROI2 bivariate distribution on day 9 and day 19 | Bivariate distribution of ROI1 and ROI2 on day 9 and day 19 |
| Figure 9C | Mann-Whitney, corrected for multiple comparisons with Benjamini-Hochberg (pymannkendall package in Python) | Test the significance of the difference in $\Delta F/F_{0peak}$ distribution between.<br>Day 1 and day 4 in cortical OBL of CLMF rule-change mice. | $\Delta F/F_{0peak}$ values in named cortical regions on each day for all mice along with their group labels. |
|  |  | day 4 and day 5 in cortical OBL of CLMF rule-change mice<br>Day 1 and Day 4 in cortical FLL of CLMF rule-change mice.<br>day 4 and day 5 in cortical FLL of CLMF rule-change mice<br>Day 1 and Day 4 in cortical V1L of CLMF rule-change mice.<br>day 4 and day 5 in cortical V1L of CLMF rule-change mice |  |
| Figure 10B, 10C | Two-way RM-ANOVA, Bonferroni corrected for multiple comparisons (Pingouin package in Python) | Test significant changes in seed pixel correlations over days and between trial and rest conditions for CLMF No-rule-change mice (Figure 10B) and for CLMF No-feedback mice (Figure 10C). | Pairwise seed pixel correlation values for all combination of cortical seed locations over days for each mouse, along with group labels. |
| Supplementary Figure 3A | Mann-Whitney U-test with Benjamini-Hochberg correction (pymannkendall package in Python) | Test the difference in the distribution of CLNF success rates between.<br>day 1 male and day 3 male.<br>day 4 male and day 4 female.<br>day 10 female and day 11 female | Normalized success rate per mouse along with their sex labels |
| Supplementary Figure 3B | Mann-Whitney U-test with Benjamini-Hochberg correction (pymannkendall package in Python) | Test the difference in the distribution of CLNF success rates between.<br>day 1 single ROI and day 3 single ROI<br>day 10 single ROI and day 10 dual ROI | Normalized success rate per mouse along with their ROI-type of the experiment. |

### CLNF rules

The task rules for CLNF experiments involved cortical activity at selected locations. These locations were standard Allen Mouse Brain Atlas (Wang et al. 2020; Chon et al. 2019) CCF coordinates. The table below lists all the task rules we tested and what they meant in the context of CLNF experiments.

**Table 3:** List of CLNF task rules with their description.

| CLNF rule name | Description |
| --- | --- |
| M1_L | $\Delta F/F_0$ activity in left hemisphere primary motor area |
| M1_R | $\Delta F/F_0$ activity in right hemisphere primary motor area |
| M2_L | $\Delta F/F_0$ activity in left hemisphere secondary motor area |
| HL_L | $\Delta F/F_0$ activity in left hemisphere sensory hindlimb area |
| RL_L | $\Delta F/F_0$ activity in left hemisphere rostralateral area of visual cortex |
| RL_R | $\Delta F/F_0$ activity in right hemisphere rostralateral area of visual cortex |
| V1_L | $\Delta F/F_0$ activity in left hemisphere primary visual area |
| BC_L – BC_R | $\Delta F/F_0$ activity in left hemisphere barrel cortex subtracted by $\Delta F/F_0$ activity in right hemisphere barrel cortex |
| HL_L – HL_R | $\Delta F/F_0$ activity in left sensory hindlimb area subtracted by $\Delta F/F_0$ activity in right hemisphere sensory hindlimb area |
| BC_L – HL_L | $\Delta F/F_0$ activity in left hemisphere barrel cortex subtracted by $\Delta F/F_0$ activity in left hemisphere sensory hindlimb area |
| M1_L – HL_L | $\Delta F/F_0$ activity in left hemisphere primary motor area subtracted by $\Delta F/F_0$ activity in left hemisphere sensory hindlimb area |
| TR_L - M2_L | $\Delta F/F_0$ activity in the left hemisphere rostral part of the temporal association area is subtracted by $\Delta F/F_0$ activity in the left hemisphere secondary motor area. |
| V1_R - V1_L | $\Delta F/F_0$ activity in the right hemisphere primary visual area subtracted by $\Delta F/F_0$ activity in the left hemisphere primary visual area. |
| M2_L - TR_L | $\Delta F/F_0$ activity in left hemisphere secondary motor area subtracted by $\Delta F/F_0$ activity in left hemisphere rostral part of temporal association area |
| HL_L - M2_L | $\Delta F/F_0$ activity in left hemisphere sensory hindlimb area subtracted by $\Delta F/F_0$ activity in left hemisphere secondary motor area |

### CLMF rules

The task rules for CLMF experiments involved points on body parts that were tracked (listed in results section). The table below lists the task rules (i.e. tracked points) we tested and what they meant in the context of CLMF experiments.

**Table 4:** List of CLMF task rules with their description.

| CLMF rule name | Description |
| --- | --- |
| FLL | Speed of left forelimb in the bottom view of the camera |
| FLR | Speed of right forelimb in the bottom view of the camera |

**Table 5:** List of mice – CLNF.

|  | MouseID | group | ROIs | initial rule | change rule | DoB | Expt. Start date | Age(days) | Sex | Reward rate |  |  |  | genotype |
| --- | --- | --- | --- | --- | --- | --- | --- | --- | --- | --- | --- | --- | --- | --- |
|  |  |  |  |  |  |  |  |  |  | Day1 | Day5 | Day11 | Day16 |  |
| 1 | BM2 | G1 | 2ROI | BC L – BC R | - | 20220122 | 20220713 | 172 | m | 0.01 | 0.8 | - | - | tta-gcamp6s |
| 2 | BM3 | G1 | 2ROI | BC L – BC R | - | 20220122 | 20220712 | 171 | m | 0.28 | 1 | - | - | tta-gcamp6s |
| 3 | BM4 | G1 | 2ROI | BC L – BC R | - | 20220122 | 20220712 | 171 | m | 0.01 | 0.83 | - | - | tta-gcamp6s |
| 4 | BG2 | G1 | 2ROI | HL L – HL R | - | 20210601 | 20211023 | 144 | m | 0.00 | 0.63 | - | - | tta-gcamp6s |
| 5 | BG3 | G1 | 2ROI | HL L – HL R | - | 20210601 | 20211023 | 144 | m | 0.00 | 0.64 | - | - | tta-gcamp6s |
| 6 | AC4 | G1 | 2ROI | HL L – HL R | - | 20210610 | 20211009 | 121 | f | 0.00 | 0.71 | - | - | tta-gcamp6s |
| 7 | TW1 | G1 | 1ROI | RL L | - | 20200729 | 20210113 | 168 | m | 0.00 | 0.35 | - | - | tta-gcamp6s |
| 8 | FM1 | G1 | 1ROI | M1 L | - | 20190915 | 20200129 | 136 | f | 0.13 | 0.66 | - | - | tta-gcamp6s |
| 9 | FM2 | G1 | 1ROI | M1 L | - | 20190915 | 20200129 | 136 | f | 0.03 | 1 | - | - | tta-gcamp6s |
| 10 | FM3 | G1 | 1ROI | M1 L | - | 20190915 | 20200129 | 136 | f | 0.02 | 1 | - | - | tta-gcamp6s |
| 11 | FM4 | G1 | 1ROI | M1 L | - | 20190915 | 20200129 | 136 | f | 0.25 | 1 | - | - | tta-gcamp6s |
| 12 | CV2 | G1 | 2ROI | BC L – HL L | - | 20190815 | 20191205 | 112 | m | 0.01 | 0.72 | - | - | tta-gcamp6s |
| 13 | CV3 | G1 | 2ROI | BC L – HL L | - | 20190815 | 20191205 | 112 | m | 0.01 | 1 | - | - | tta-gcamp6s |
| 14 | DU1 | G1 | 2ROI | M1 L – HL L | - | 20190719 | 20191205 | 139 | f | 0.01 | 1 | - | - | tta-gcamp6s |
| 15 | DU3 | G1 | 2ROI | M1 L – HL L | - | 20190719 | 20191205 | 139 | f | 0.02 | 0.39 | - | - | tta-gcamp6s |
| 16 | BZ1 | G1 | 2ROI | TR L – M2 L | - | 20190406 | 20190806 | 122 | m | 0.24 | 0.29 | - | - | tta-gcamp6s |
| 17 | BZ2 | G1 | 2ROI | TR L – M2 L | - | 20190406 | 20190806 | 122 | m | 0.02 | 0.32 | - | - | tta-gcamp6s |
| 18 | DU1 | G1 | 2ROI | V1 R – V1 L | - | 20190217 | 20190807 | 171 | f | 0.00 | 1 | - | - | tta-gcamp6s |
| 19 | DU2 | G1 | 2ROI | V1 R – V1 L | - | 20190217 | 20190807 | 171 | f | 0.00 | 0.46 | - | - | tta-gcamp6s |
| 20 | BZ1 | G1 | 2ROI | M2 L – TR L | - | 20181230 | 20190516 | 137 | m | 0.19 | 0.19 | - | - | tta-gcamp6s |
| 21 | BZ2 | G1 | 2ROI | M2 L – TR L | - | 20181230 | 20190516 | 137 | m | 0.02 | 0.71 | - | - | tta-gcamp6s |
| 22 | BU2 | G1 | 2ROI | M2 L – TR L | - | 20181219 | 20190517 | 149 | m | 0.20 | 1 | - | - | tta-gcamp6s |
| 23 | BU3 | G1 | 2ROI | M2 L – TR L | - | 20181219 | 20190517 | 149 | m | 0.01 | 0.61 | - | - | tta-gcamp6s |
| 24 | BM1 | G2 | 2ROI | BC L – BC R | RFLL – CFLL (Day11) | 20220122 | 20220712 | 171 | m | 0.00 | 0.63 | 0.03 | 0.52 | tta-gcamp6s |
| 25 | AC1 | G2 | 2ROI | BC L – BC R | HL L – HL R (Day11) | 20210610 | 20211009 | 121 | f | 0.27 | 0.41 | 0.12 | 1 | tta-gcamp6s |
| 26 | AC2 | G2 | 2ROI | BC L – BC R | HL L – HL R (Day11) | 20210610 | 20211009 | 121 | f | 0.00 | 1 | 0.29 | 0.96 | tta-gcamp6s |
| 27 | TW1 | G2 | 1ROI | RL L | HL L (Day11) | 20200802 | 20210113 | 129 | m | 0.01 | 0.58 | 0.00 | 0.71 | tta-gcamp6s |
| 28 | TW2 | G2 | 1ROI | RL L | PM L (Day11) | 20200802 | 20210113 | 129 | m | 0.02 | 0.28 | 0.32 | 0.90 | tta-gcamp6s |
| 29 | TW22 | G2 | 1ROI | RL R | RL L (Day11) | 20200730 | 20201119 | 112 | m | 0.10 | 0.48 | 0.35 | 0.60 | tta-gcamp6s |
| 30 | TW11 | G2 | 1ROI | M2 L | RL L (Day11) | 20200730 | 20201119 | 112 | m | 0.05 | 0.32 | 0.10 | 0.64 | tta-gcamp6s |
| 31 | TW33 | G2 | 1ROI | M1 R | RL R (Day11) | 20200730 | 20201119 | 112 | m | 0.02 | 0.41 | 0.40 | 0.60 | tta-gcamp6s |
| 32 | FM3 | G2 | 1ROI | V1 L | M1_R (Day11) | 20190917 | 20200129 | 134 | f | 0.08 | 0.62 | 0.03 | 0.37 | tta-gcamp6s |
| 33 | FM1 | G2 | 1ROI | M1 L | V1 L (Day11) | 20190917 | 20200129 | 134 | f | 0.00 | 0.65 | 0.02 | 0.61 | tta-gcamp6s |
| 34 | FM2 | G2 | 1ROI | M1 L | M1_R (Day11) | 20190917 | 20200129 | 134 | f | 0.10 | 1 | 0.14 | 1 | tta-gcamp6s |
| 35 | FM3 | G2 | 1ROI | M1 L | V1 L (Day11) | 20190917 | 20200129 | 134 | f | 0.03 | 0.63 | 0.20 | 0.46 | tta-gcamp6s |
| 36 | FM4 | G2 | 1ROI | M1 L | V1 L (Day11) | 20190917 | 20200129 | 134 | f | 0.01 | 0.31 | 0.00 | 0.62 | tta-gcamp6s |
| 37 | DR2 | G2 | 2ROI | M2 L – TR L | HL L – M2 L (Day11) | 20190401 | 20190806 | 127 | m | 0.09 | 1 | 0.01 | 1 | tta-gcamp6s |
| 38 | DR3 | G2 | 2ROI | M2 L – TR L | HL L – M2 L (Day11) | 20190401 | 20190806 | 127 | m | 0.03 | 1 | 0.06 | 0.74 | tta-gcamp6s |
| 39 | DR4 | G2 | 2ROI | M2 L – TR L | HL L – M2 L (Day11) | 20190401 | 20190806 | 127 | m | 0.00 | 0.64 | 0.02 | 1 | tta-gcamp6s |
| 40 | DU3 | G2 | 2ROI | M2 L – TR L | V1_R – V1_L (Day11) | 20190408 | 20190807 | 121 | f | 0.13 | 0.36 | 0.10 | 1 | tta-gcamp6s |

**Table 6:** List of mice – CLMF.

|  | MouseID | Group | initial_rule | change_rule | DoB | Exp. start | Age(days) | Sex | Reward rate |  |  |  | genotype |
| --- | --- | --- | --- | --- | --- | --- | --- | --- | --- | --- | --- | --- | --- |
|  |  |  |  |  |  |  |  |  | Day1 | Day4 | Day5 | Day10 |  |
| 1 | FW2 | Rule change | FLL | FLR (Day5) | 20230209 | 20230713 | 154 | f | 0.30 | 0.60 | 0.21 | 0.95 | Ai94 |
| 2 | FW3 | Rule change | FLL | FLR (Day5) | 20230209 | 20230718 | 159 | f | 0.09 | 0.65 | 0.00 | 0.92 | Ai94 |
| 3 | FW22 | Rule change | FLL | FLR (Day5) | 20230209 | 20230719 | 160 | f | 0.25 | 0.70 | 0.44 | 1 | Ai94 |
| 4 | GT3 | Rule change | FLL | FLR (Day5) | 20230219 | 20230715 | 146 | f | 0.11 | 0.93 | 0.63 | 0.85 | Ai94 |
| 5 | GT33 | Rule change | FLL | FLR (Day5) | 20230219 | 20230719 | 150 | f | 0.12 | 0.78 | 0.00 | 0.87 | Ai94 |
| 6 | HA2 | Rule change | FLL | FLR (Day5) | 20230225 | 20230719 | 144 | m | 0.12 | 0.72 | 0.28 | 0.9 | Ai94 |
| 7 | GER2 | Rule change | FLL | FLR (Day5) | 20230823 | 20231228 | 127 | m | 0.34 | 0.95 | 0.50 | 0.83 | Ai94 |
| 8 | HYL3 | Rule change | FLL | FLR (Day5) | 20230827 | 20240103 | 129 | m | 0.33 | 0.93 | 0.60 | 0.91 | Ai94 |
| 9 | BR1 | No rule change | FLL | - | 20230729 | 20231201 | 125 | m | 0.11 | 0.58 | 0.97 | 1 | Ai94 |
| 10 | BR2 | No rule change | FLL | - | 20230729 | 20231201 | 125 | m | 0.38 | 1 | 0.84 | 0.82 | Ai94 |
| 11 | GER1 | No rule change | FLL | - | 20230811 | 20231228 | 139 | f | 0.35 | 0.88 | 0.84 | 0.82 | Ai94 |
| 12 | HYL2 | No rule change | FLL | - | 20240103 | 20240103 | 121 | f | 0.33 | 0.93 | 0.88 | 0.93 | Ai94 |
| 13 | GIL2 | No feedback | FLL | - | 20231219 | 20231219 | 112 | m | 0.09 | 0.52 | 0.28 | 0.95 | Ai94 |
| 14 | GIL3 | No feedback | FLL | - | 20231219 | 20231219 | 135 | f | 0.23 | 0.53 | 0.57 | 0.51 | Ai94 |
| 15 | GIL4 | No feedback | FLL | - | 20231219 | 20231219 | 135 | f | 0.19 | 0.63 | 0.60 | 0.69 | Ai94 |
| 16 | GER3 | No feedback | FLL | - | 20230811 | 20231219 | 130 | m | 0.08 | 0.48 | 0.43 | 0.75 | Ai94 |

## Supplementary

**Animation 1:**
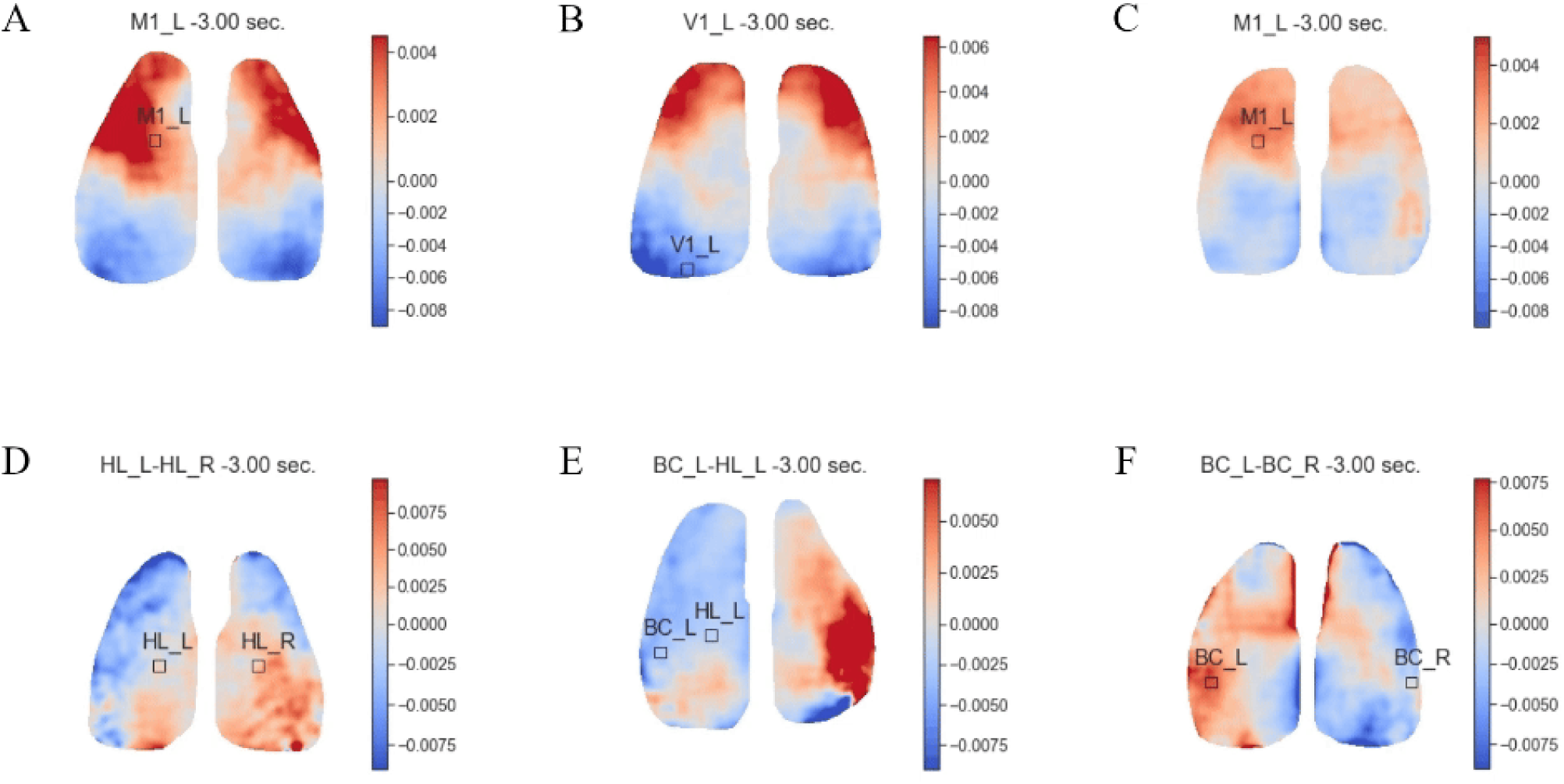
CLNF success centered trial averages under various task rules. A) Success centered ΔF/F_0_ cortical dynamics in 1ROI experiment with target task rule as M1_L B) Success centered ΔF/F_0_ cortical dynamics in 1ROI experiment with target task rule as V1_L C) Success centered ΔF/F_0_ cortical dynamics in 1ROI experiment with target task rule as M1_L D) Success centered ΔF/F_0_ cortical dynamics in 2ROI experiment with target task rule as HL_L - HL_R E) Success centered ΔF/F_0_ cortical dynamics in 2ROI experiment with target task rule as BC_L - HL_L F) Success centered ΔF/F_0_ cortical dynamics in 2ROI experiment with target task rule as BC_L - BC_R

**Animation 2:**
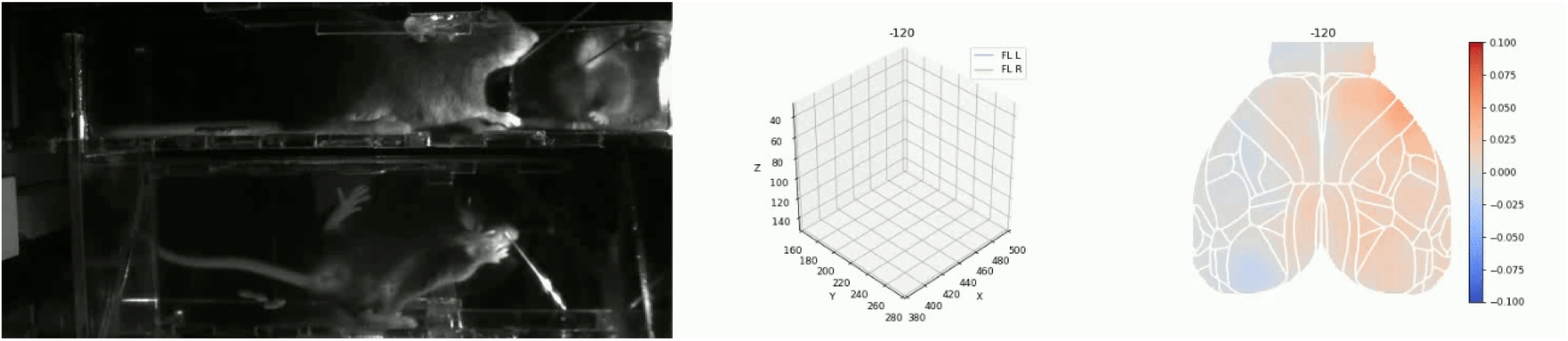
CLMF single trial with FLL_bottom as control point. Behavior video on the left panel, two of the tracked body parts (FLL_bottom, FLR_bottom) at the center, and corresponding cortical widefield ΔF/F_0_ activity of a trial with target control point as FLL_bottom (left forelimb in camera bottom view) on the right panel.

**Animation 3:**
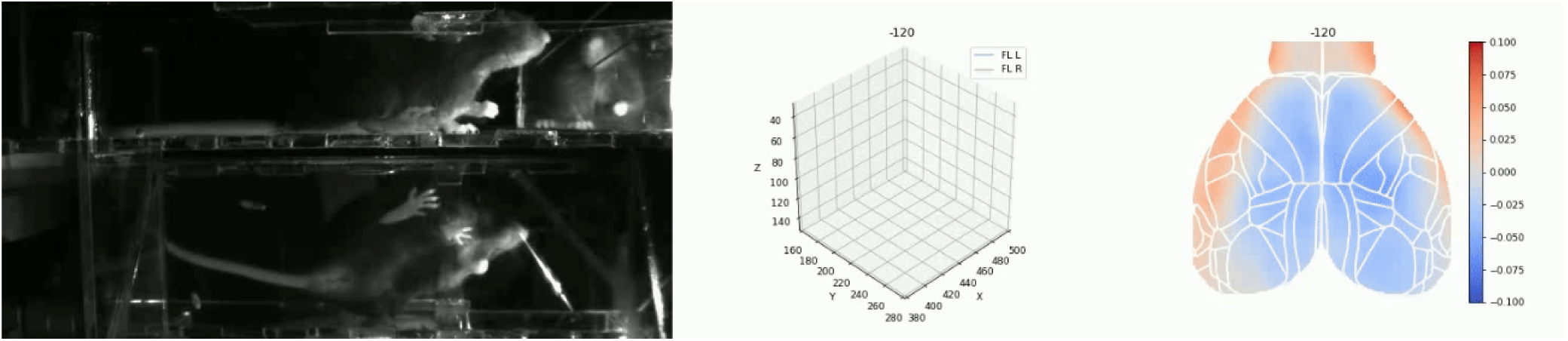
CLMF single trial with FLR_bottom as control point. Behavior video on the left panel, two of the tracked body parts (FLL_bottom, FLR_bottom) at the center, and corresponding cortical widefield ΔF/F_0_ activity of a trial with target control point as FLR_bottom (right forelimb in camera bottom view) in the same mouse shown in Animation 2 on the right panel.

**Supplementary Figure 1:**
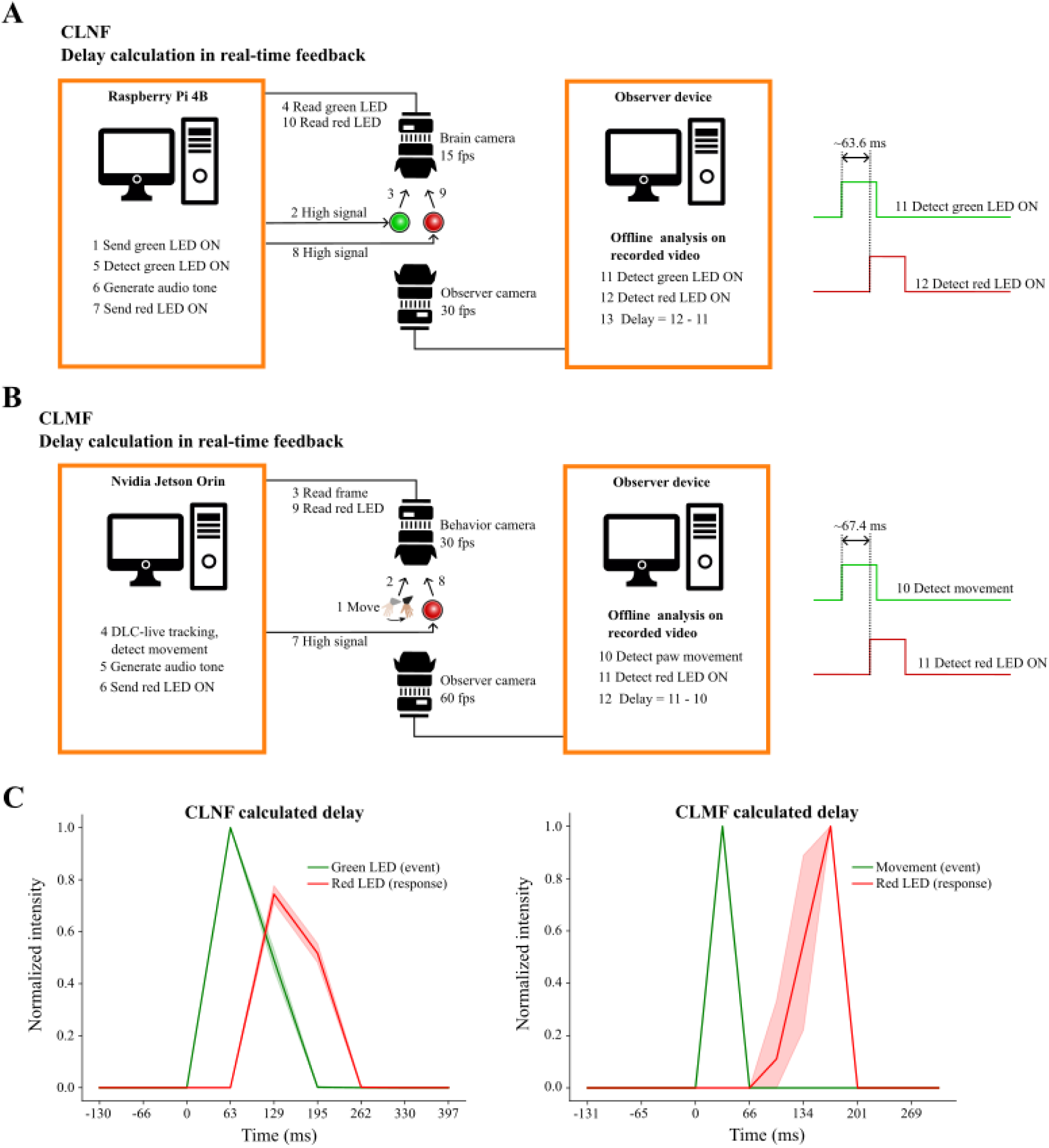
Method for calculating feedback and reward delay in CLNF and CLMF setups. Two devices (device running closed-loop system, an observer device that was just recording the scene), each connected to a camera with overlapping field of view were employed. A) Setup for calculating feedback delay in CLNF experiments deployed on Raspberry Pi. The difference between the onset of the green LED and red LED ON events is the feedback delay. 1) The Raspberry Pi generates a HIGH signal on the GPIO pin connected to the green LED. 2) The HIGH signal is received at the anode of the green LED, causing it to glow. 3) Light emitted by the green LED is received by the brain camera sensor. 4) An image capturing the green emitted light is captured and sent to the Raspberry Pi. 5) The CLNF program processes the image and detects changes in brightness. 6) The CLNF program maps the brightness change to an audio frequency and generates an audio tone output at that frequency. 7) The CLNF program sends a HIGH signal to the GPIO pin connected to the red LED. 8) The HIGH signal is received at the anode of the red LED, causing it to glow. 9) Light emitted by the red LED is received by the sensor of the brain camera. 10) An image frame capturing the red LED light is captured and sent to the Raspberry Pi. 11) During the previous steps, an observer camera records the green and red LEDs. In offline analysis, detect the brightness change in the green LED. 12) Detect the brightness change in the red LED. 13) Calculate the response delay as the time difference between the green and red LED brightness change events. B) Setup for calculating feedback delay in CLMF experiments deployed on Nvidia Jetson Orin. The difference between the onset of movement and the red LED ON events is the feedback delay. 1) The head-fixed mouse in the CLMF apparatus moved the control point crossing the speed threshold. 2) Behavior is captured by the behavior camera and observer camera sensors. 3) The captured image is sent to Nvidia Jetson running the CLMF program. 4) The image frame is analyzed, and tracked points are detected. Calculate the speed of the control point. 5) Map the control point speed to an audio frequency and generate audio tone output at the frequency. 6) Send a HIGH signal to the GPIO pin connected to the red LED. 7) The HIGH signal is received at the anode of the red LED. 8) Bright light from the red LED reaches the behavior camera and observer camera sensors. 9) The captured image is sent to Nvidia Jetson and the observer device. 10) In offline analysis, detect the control point movement event. 11) Detect the brightness change event in the red LED. 12) Calculate the response delay as the time difference between the control point movement event and the red LED brightness change event. C) Offline analysis to calculate response time using data collected in an experiment. Left: Brightness change events of green LED and red LED in CLNF setup, data represents 100 trials. Right: Control-point movement event and brightness change event of red LED in CLMF setup, data represents 30 trials.

**Supplementary Figure 2:**
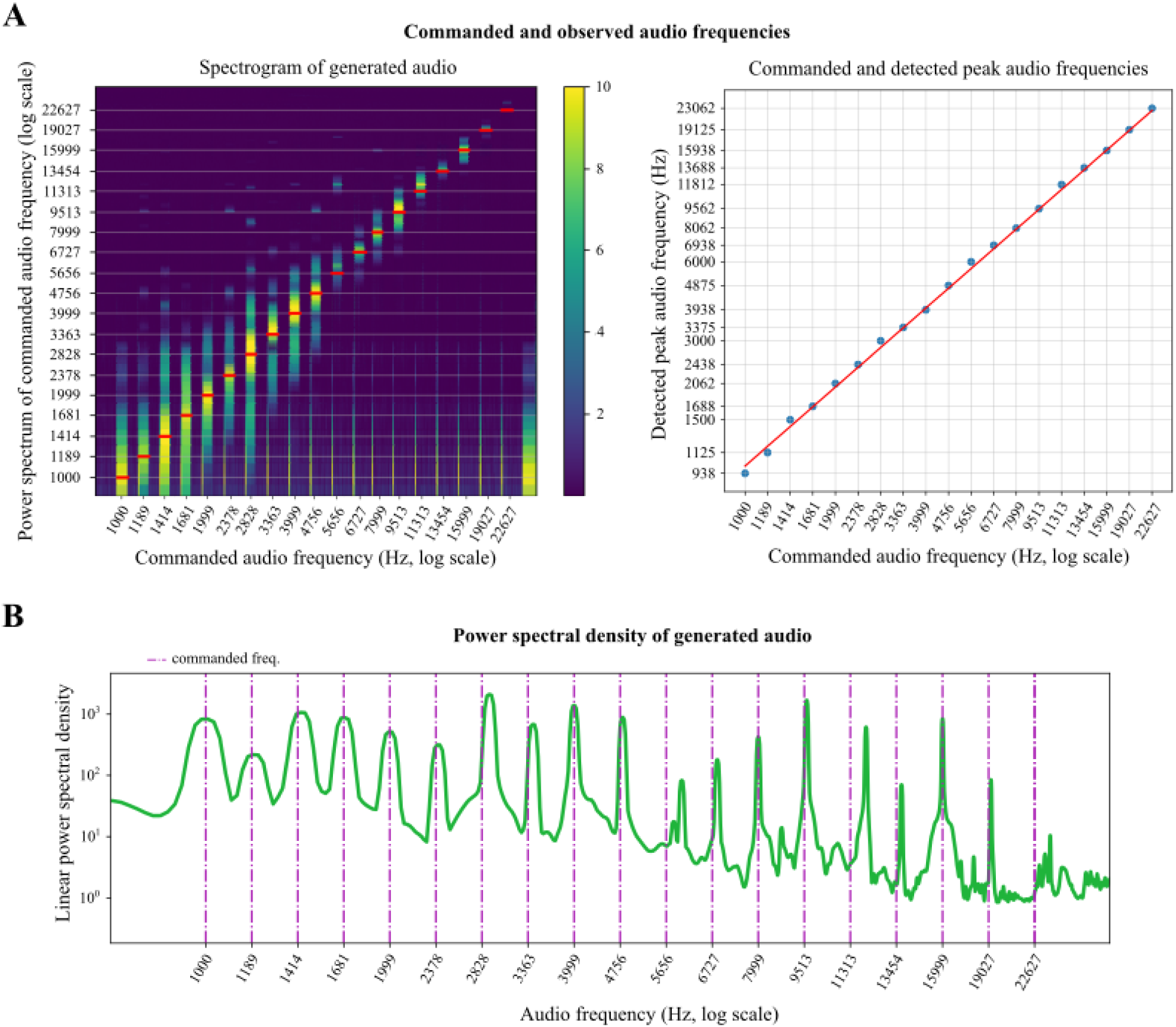
Verification of generated audio tone frequencies. A) Left: Spectrogram of an audio recorded in a session where graded audio tone frequencies were generated sequentially from 1 kHz to 22.627 kHz using the CLNF program. Desired frequency values are shown on the x-axis and expected frequencies are marked in red on the y-axis. The frequency values increase exponentially, and therefore the power bands shrink as frequency increases. Right: Peak powers detected compared to the intended frequency, using the spectrogram on the left. B) The power spectral density of the audio recording used is A. The width of the peaks becomes narrower as the frequency increases since the x-axis is on a log scale.

**Supplementary Figure 3:**
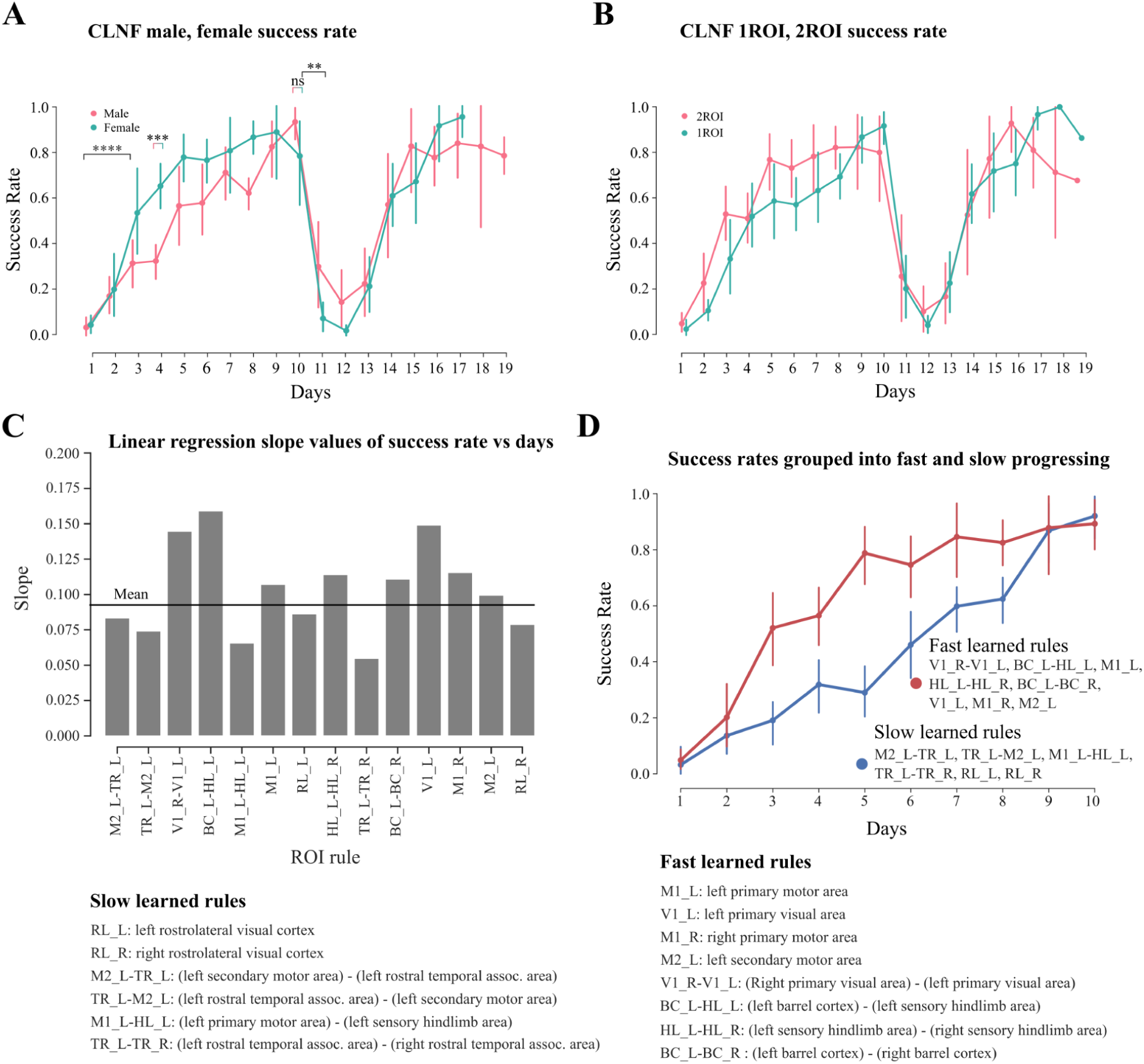
Learning progression was similar in male and female mice, and even similar between 1ROI and 2ROI experiments. There were some ROI rules that were faster to learn than others. A) The success rates over days in male and female mice are not significantly different. Data from all mice in the No-rule-change and Rule-change groups were separated based on their sex (male, female). For days 1-10,the total number of males was 14 and the total number of females was 9; for days 11-19,the total number of males was 9 and the total number of females was 9. B) Success rate over days based on 1ROI and 2ROI experiments. Data from all mice in the No-rule-change and Rule-change groups were segregated based on the ROI type (1ROI or 2ROI). From Day 1 to Day 10,the total number of mice with 1ROI experiments was 5, and with 2ROI it was 18. From Day 11 to Day 19,the total number of mice with 1ROI was 10, and with 2ROI it was 7. C) Slope values of linear regression line on the success rate over days for different ROI-rules. D) ROI rules with slope values greater than or equal to the mean (0.095) were combined in the red group, and those with slope values lower than the mean were combined in the blue group. At a broad level, we can observe that ROI rules in the red group were faster than those in the blue group.

**Supplementary Figure 4:**
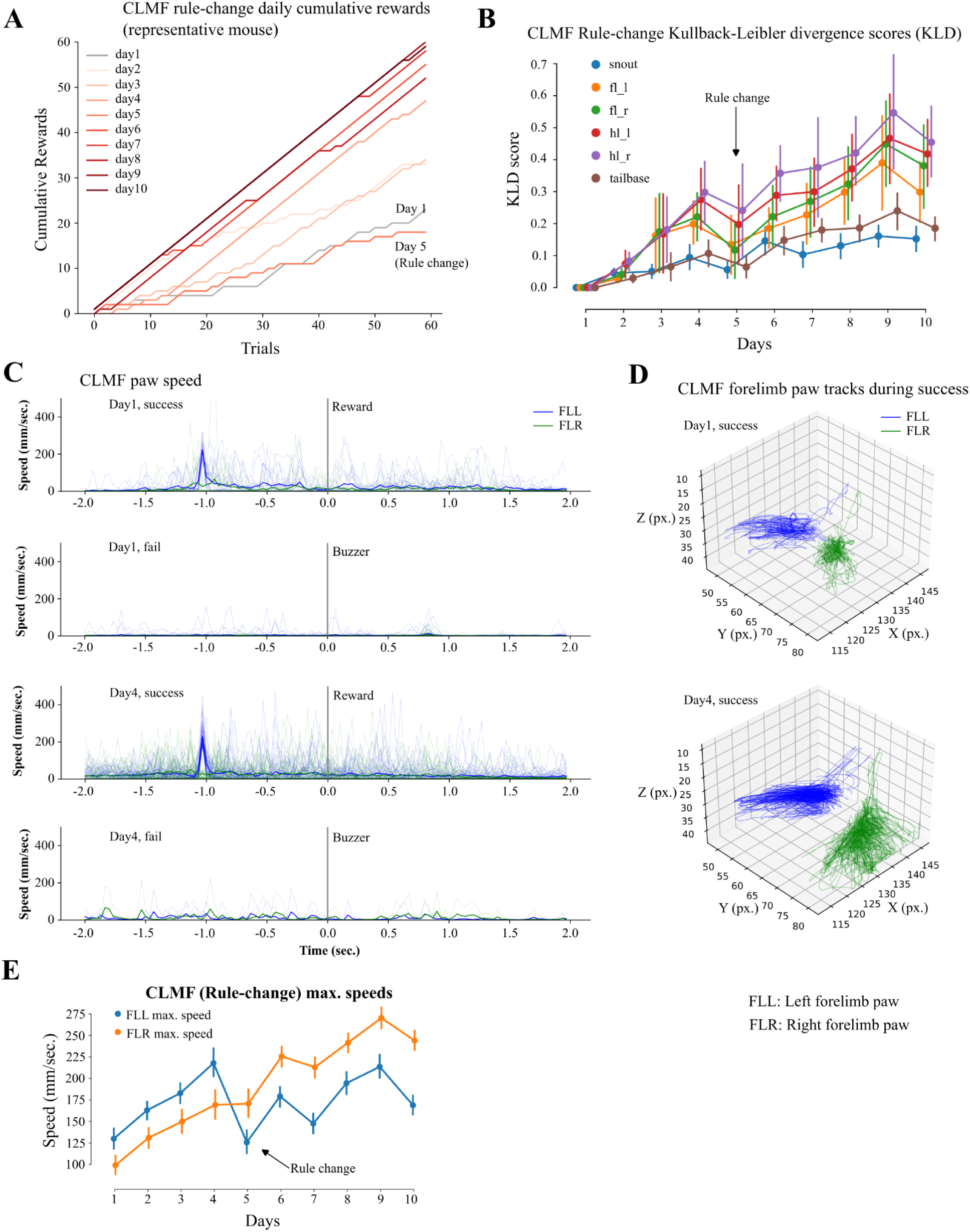
CLMF behavior traces. A) Cumulative rewards show performance during the whole session from day 1 to day 10, including a perturbation on day 5 where cumulative rewards drops, in a CLMF rule-change mouse. B) Kullback-Leibler divergence scores of speed distributions of different tracked points (snout, left forelimb, right forelimb, left hind limb, right hind limb, and tail base) on the mouse body. Kernel density estimates (KDE) were derived for these tracked points on each day, and divergence scores compared to day 1 were calculated. C) Reward centered left (in blue) and right (in green) forelimb speeds, top row (day1) and third row (day4). Trial end centered left (in blue) and right (in green) forelimb speeds, second (day1) and fourth row (day4). The target control-point during training was left forelimb. D) Left (in blue) and right (in green) forelimb tracks on day 1 (top) and day 4 (bottom) with the control point being the left forelimb.

**Supplementary Figure 5:**
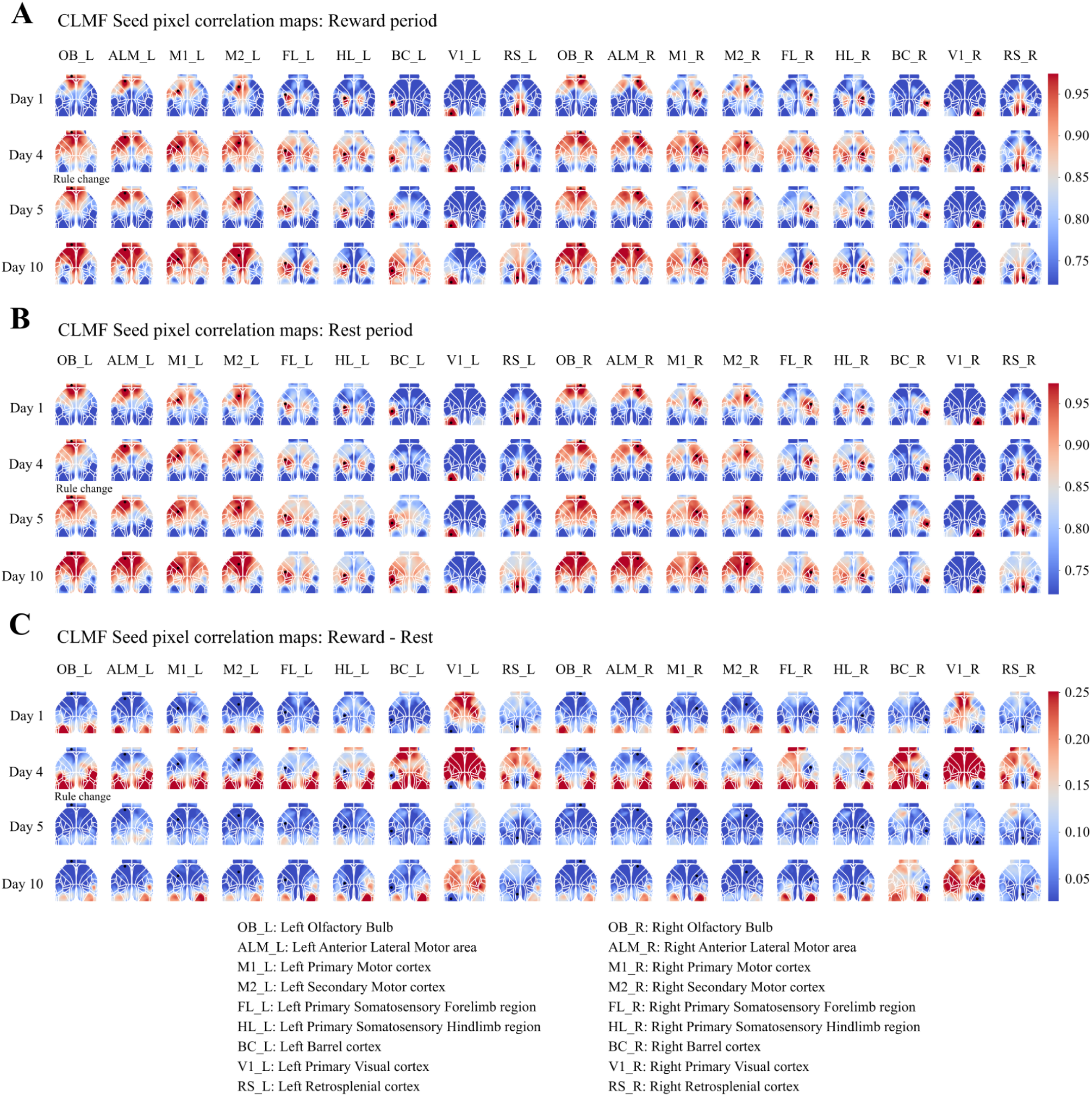
CLMF Seed pixel correlation maps during rewarded trial, rest periods, and their difference. A) Seed pixel correlation maps of labeled seed pixels on day 1, day 4, day 5 (rule change), and day 10 during rewarded trials (excluding the reward consumption period). Seed pixel locations are indicated by black points on the brain map. B) Seed pixel correlation maps of labeled seed pixels on day 1, day 4, day 5 (rule change), and day 10 during rest periods (excluding the reward consumption period). Seed pixel locations are indicated by black points on the brain map. C) Difference of seed pixel correlation maps of labeled seed pixels during rewarded trials and rest periods (A - B) on day 1, day 4, day 5 (rule change), and day 10. Seed pixel locations are indicated by black points on the brain map. There is a distinct increase in correlation during rewarded trials in cortical regions such as V1 and RS across all seed pixel maps.

**Supplementary Figure 6:**
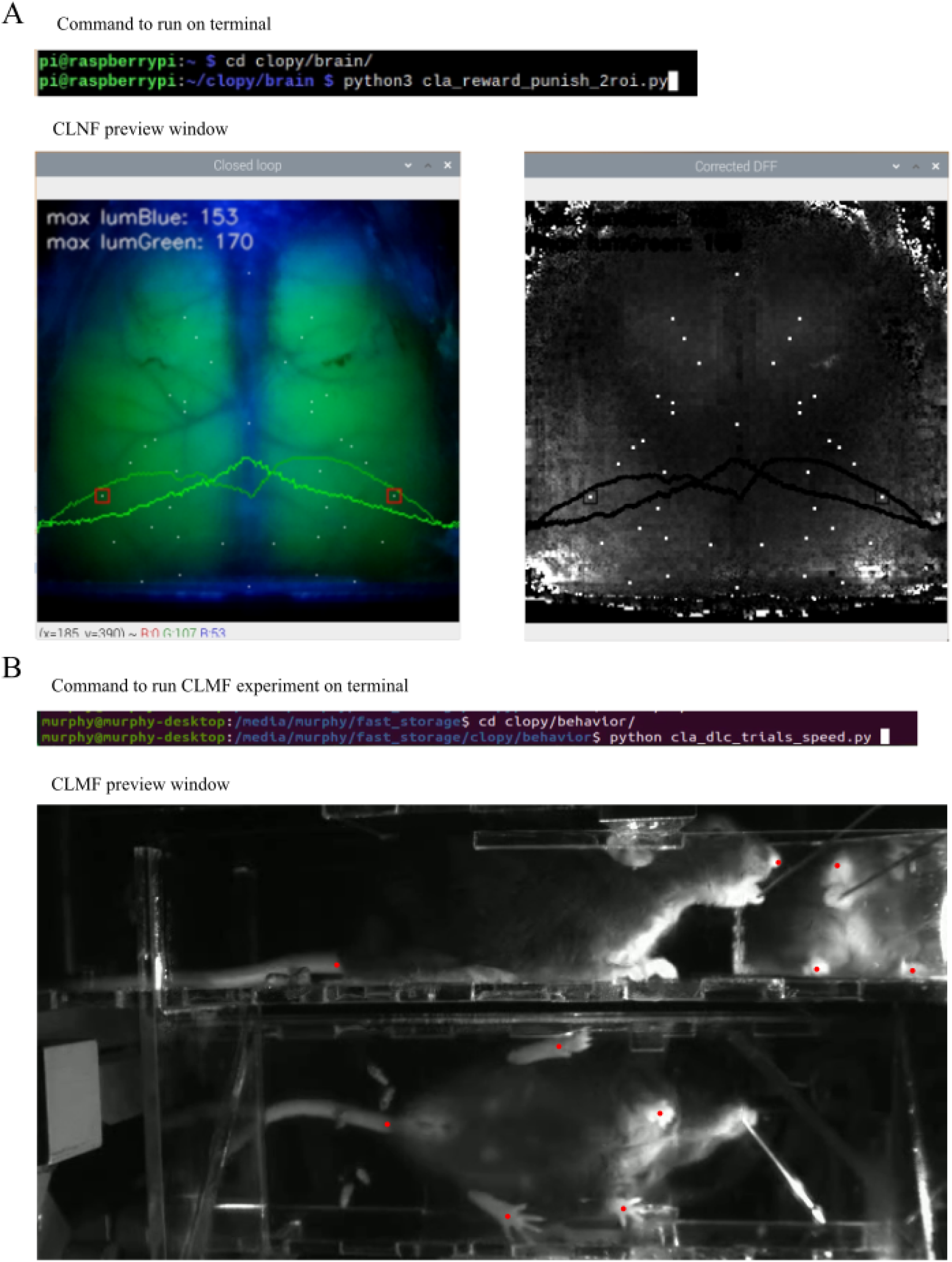
CLNF, CLMF command line operation and preview steps. A) Top: Terminal commands to launch a CLNF experiment. Bottom left: Showing widefield calcium activity (green) and reflectance (blue) signals as a composite RGB image. Overlaid text indicates maximum green and blue intensities in the middle row of the image; bottom curves show intensity in the green channel (faded green) and blue channel (bright green) along the middle row of pixels. Two red squares indicate the ROIs in the 2ROI experiment. White dots are the registered centers of the cortical locations specified in the config.ini file. Bottom right: Showing the corrected ΔF/F_0_ map of the image on its left in real-time. The overlaid artifacts in the form of text and lines are there since we draw those text and lines on the captured image for preview. These artifacts are not present when a session starts. B) Top: Terminal commands to launch a CLMF experiment. Bottom: The preview window shows a live view with tracked points overlaid on the live view, indicated in red. To start the experiment, press ‘Esc’ on the keyboard, type the mouse ID in the prompt, and then press the Enter button to start the experiment.

**Supplementary Figure 7:**
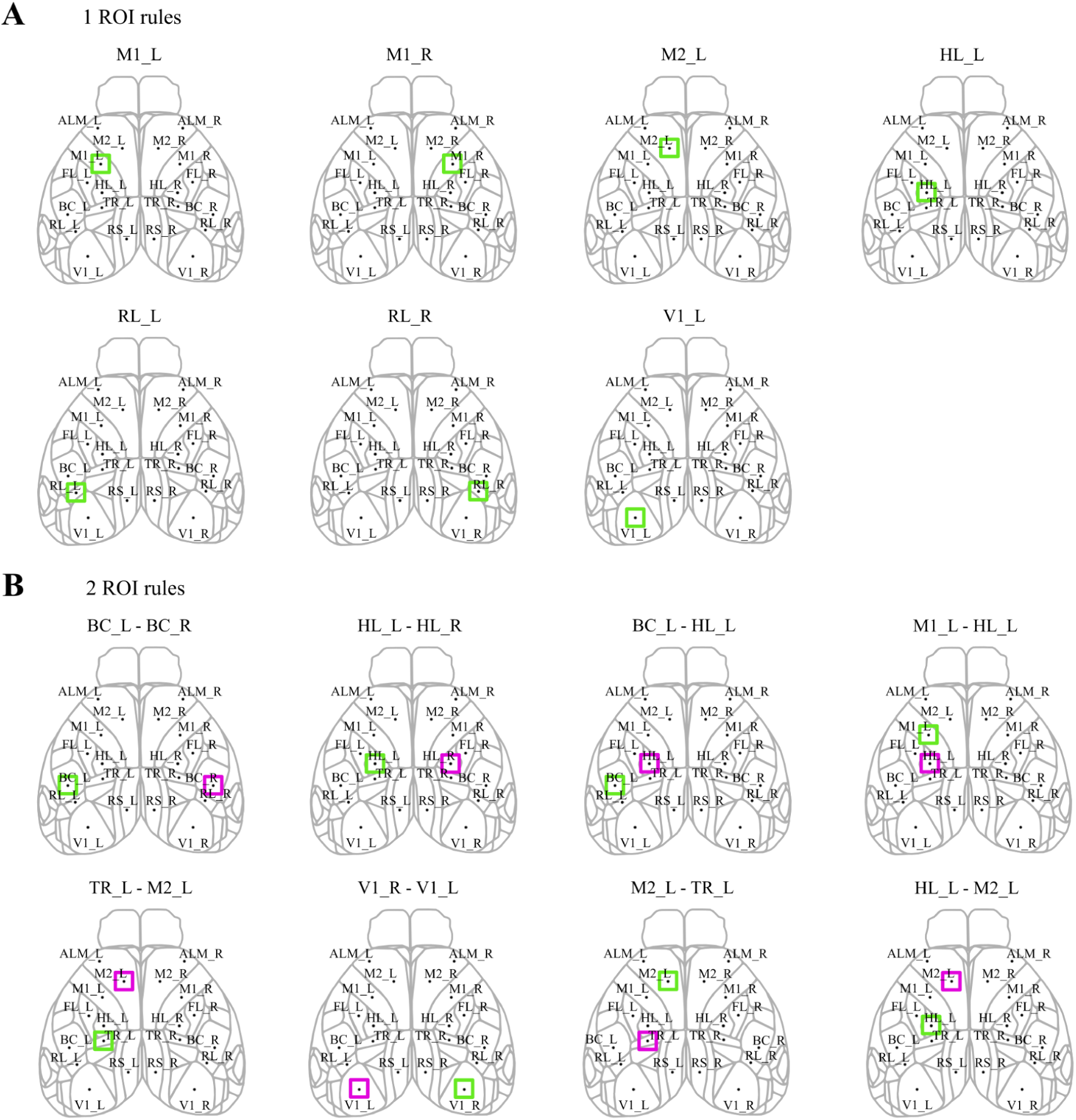
Visual representation of 1 ROI and 2 ROI rules in CLNF experiment. A) ROIs overlaid on mouse dorsal cortical map during 1 ROI experiments. B) ROIs overlaid on mouse dorsal cortical map during 2 ROI experiments.

**Supplementary Figure 8:**
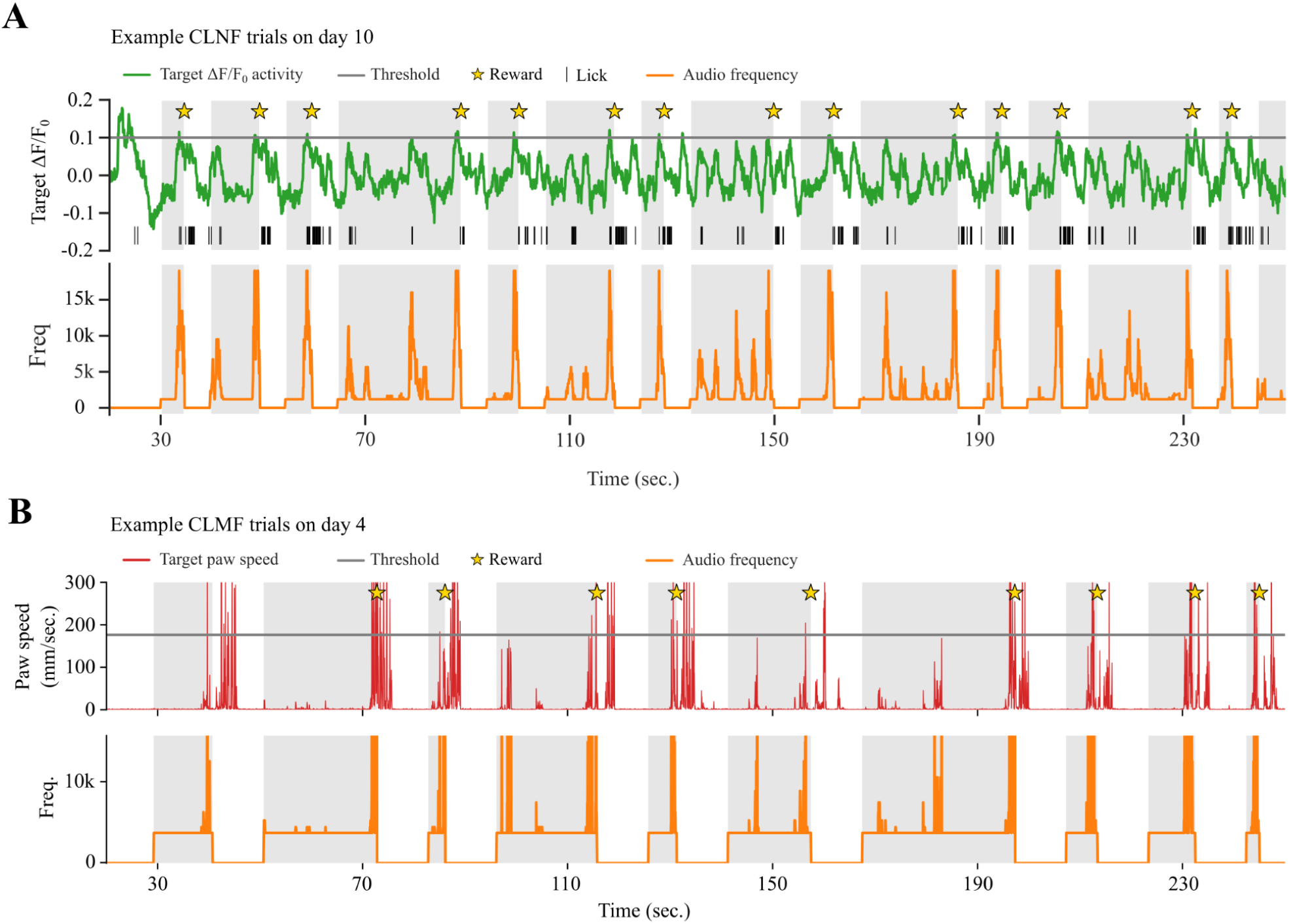
Example trials along with audio feedback during CLNF and CLMF training. A) Target cortical activity in Green, along with horizontal threshold line, Yellow stars as rewards, Black ticks showing licks, and Orange trace showing the audio frequency. B) Target paw speed in Red, along with horizontal threshold like, Yellow stars as rewards, and Orange trace showing the audio frequency.

