## Supplementary figures and images for "Real-Time Closed-Loop Feedback System For Mouse Mesoscale Cortical Signal And Movement Control: CLoPy"

### supplemental animation 1 closed-loop neurofeedback

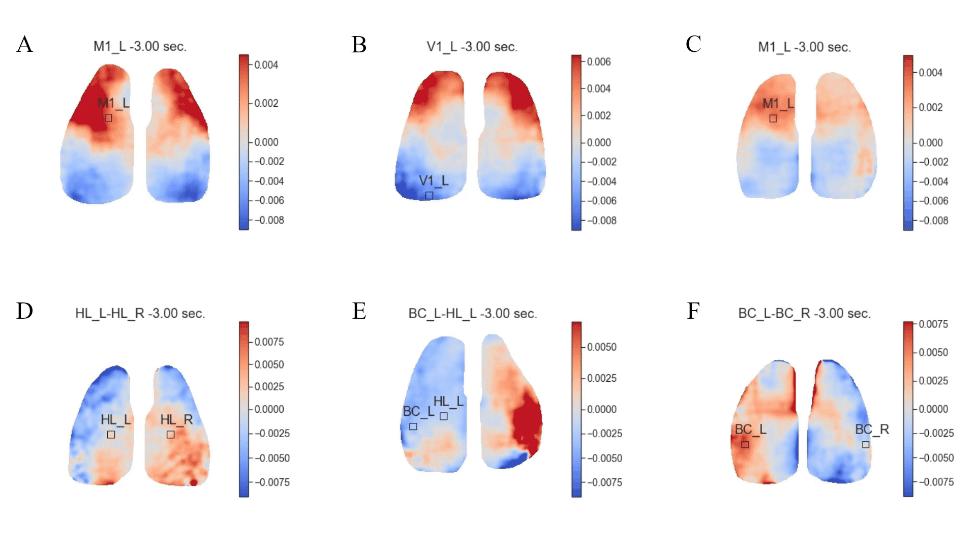

### supplemental animation 2 closed-loop movement feedback forelimb left

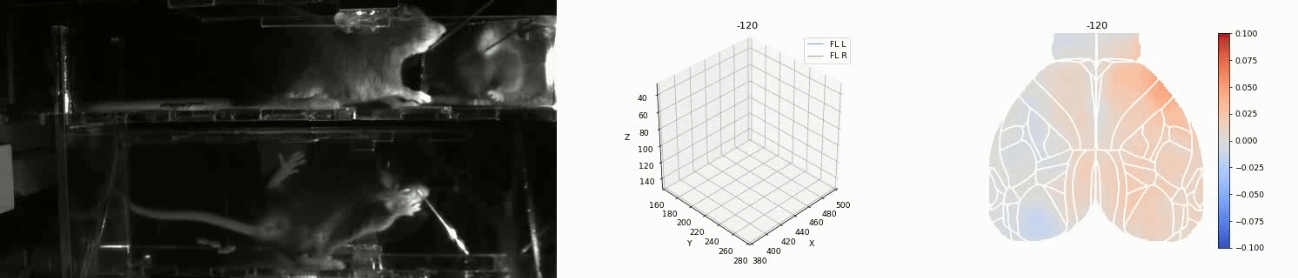

### supplemental animation 3 closed-loop movement feedback forelimb right

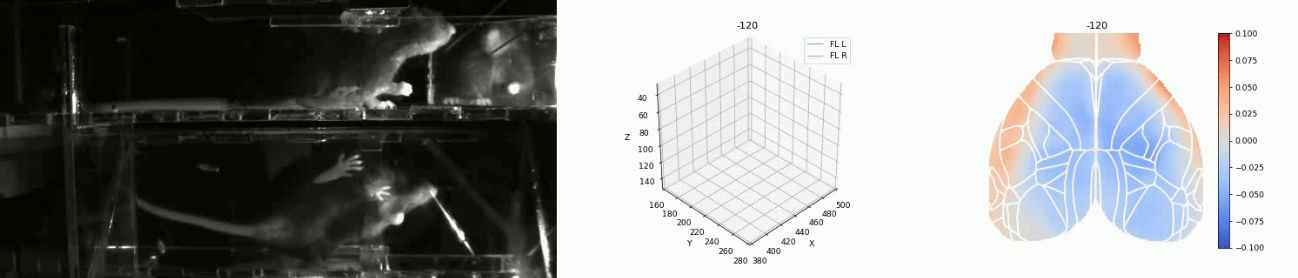
